# Activation of TnSmu1, an integrative and conjugative element, by an ImmR-like transcriptional regulator in *Streptococcus mutans*

**DOI:** 10.1101/2022.05.11.491493

**Authors:** Shawn King, Allison Quick, Kalee King, Alejandro R. Walker, Robert C. Shields

## Abstract

Integrative and conjugative elements (ICEs) are chromosomally encoded mobile genetic elements that can transfer DNA between bacterial strains. Recently, as part of efforts to determine hypothetical gene functions, we have discovered an important regulatory module encoded on an ICE known as TnSmu1 on the *Streptococcus mutans* chromosome. The regulatory module consists of a cI-like repressor with a helix-turn-helix DNA binding domain *immR*_Smu_ (immunity repressor) and a metalloprotease *immA*_Smu_ (anti-repressor). It is not possible to create an in-frame deletion mutant of *immR*_Smu_ and repression of *immR*_Smu_ with CRISPRi causes substantial cell defects. We used a bypass of essentiality (BoE) screen to discover genes that allow deletion of the regulatory module. This revealed that conjugation genes, located within TnSmu1, can restore the viability of an *immR*_Smu_ mutant. Deletion of *immR*_Smu_ also leads to production of a circular intermediate form of TnSmu1, that is also inducible by the genotoxic agent mitomycin C. To gain further insights into potential regulation of TnSmu1 by ImmR_Smu_ and broader effects on *S. mutans* UA159 physiology we used CRISPRi and RNA-seq. Strongly induced genes included all the TnSmu1 mobile element, genes involved in amino acid metabolism, transport systems, and a Type I-C CRISPR-Cas system. Lastly, bioinformatic analysis shows that the TnSmu1 mobile element and its associated genes are well distributed across *S. mutans* isolates. Taken together, our results show that activation of TnSmu1 is controlled by the *immRA*_Smu_ module, and that activation is deleterious to *S. mutans*, highlighting the complex interplay between mobile elements and their host.

## INTRODUCTION

*Streptococcus mutans* is a Gram-positive bacterium that colonizes the human oral cavity [1]. Like many streptococci, *S. mutans* can cause disease when environmental conditions are favorable. Poor oral hygiene combined with frequent ingestion of simple sugars creates an environment in which *S. mutans* can cause dental caries (tooth decay) [2, 3]. If this disease is allowed to progress it causes the breakdown of teeth, with symptoms that include pain, difficulty eating and tooth loss [2]. Due to its widespread presence in the oral cavity and resulting disease burden it is necessary to identify targets for improved therapeutics and understand processes essential for the pathogen. With this goal in mind, we have recently applied transposon sequencing (Tn-seq) and CRISPR interference (CRISPRi) and have identified >200 *S. mutans* essential genes that are required for the viability of the organism [4, 5]. For *S. mutans*, most essential genes can be broadly sorted into three categories: processing of genetic information, energy production, and maintenance of the cell envelope [4].

It was during Tn-seq/CRISPRi experiments that we discovered an essential gene, SMU_218, which is annotated as a transcriptional regulator, and is designated *immR*_Smu_ (immunity repressor) from here forwards. *immR*_Smu_ resides in a two-gene operon with SMU_219/*immA*_Smu_ (anti-repressor). Probing *immR*_Smu_ and *immA*_Smu_ with SMART (Simple Modular Architecture Research Tool) [6], we found that the N-terminus of *immR*_Smu_ contains a cI-like repressor of phage λ DNA-binding domain, often found in streptococcal phages. *immA*_Smu_ contains a putative ImmA/IrrE family metallo-endopeptidase domain, based on possession of a conserved metalloprotease zinc-binding motif, HEXXH [7]. In addition, the toxin-antitoxin database RASTA [8] annotates this two-gene operon as a putative toxin-antitoxin module. We found this interesting as the antitoxin component (*e.g. immR*_Smu_) of toxin-antitoxin modules are often essential, because they prevent the accumulation of the toxin protein. While we cannot exclude the hypothesis that this two-gene operon is a toxin-antitoxin module, exhaustive literature searches provide evidence for other functions of these genes. First, model λ cI repressors have an N-terminal domain that binds DNA and a 3’ domain that functions in cI autoproteolysis, in conjunction with RecA [9]. In *S. mutans*, it is possible that these functions are separated into two genes, *immR*_Smu_ and *immA*_Smu_. Next, cI-like repressors regulate prophage induction in bacteria but they have also been shown to regulate integrative conjugative element (ICE) expression [10, 11]. Notably, *immR*_Smu_ and *immA*_Smu_ reside in the flanking region of the putative ICE designated as TnSmu1. This large region of DNA (23 kilobases) contains predicted conjugation genes, as well as many hypothetical proteins, and is found in several strains of *S. mutans* [12, 13]. Very little is known about the activity or the mobility of this element, but it shares similarities with *ICESt1*, ICE*St3*, Tn*916* and *ICEBs1*. Although the regulatory regions of these different ICEs are substantially re-arranged compared to TnSmu1, *ICESt1* and *ICEBs1* contain cI-like repressors that repress ICE expression [10, 11, 14]. Therefore, we have formulated two major hypotheses explaining the essential nature of *immR*_Smu_: 1) *immR*_Smu_ is an antitoxin in a toxin-antitoxin module or 2) *immR*_Smu_ is involved in the activation of TnSmu1. For the first hypothesis CRISPR-mediated knockdown of *immR*_Smu_ would lead to over-accumulation of *immA*_Smu_, which would then act as a toxin. For the second hypothesis, loss of the repressor would lead to activation of TnSmu1 with loss of the element from the cell making *immR*_Smu_ appear ‘essential’.

Here we investigate both of our hypotheses for explaining the essentiality of *immR*_Smu_. Using transposon mutagenesis, genome sequencing, RNA-seq, bioinformatics and standard molecular microbiology techniques we find that *immR*_Smu_ is very likely to be a repressor of the ICE TnSmu1 and is related to the *ICEBs1* immunity repressor *immR*. Removal of *immR*_Smu_ repression causes upregulation of the TnSmu1 element leading to excision from the genome and formation of a circular intermediate. We found that TnSmu1 is activated by DNA damage and we posit that this occurs via ImmA_Smu_-mediated cleavage of the ImmR_Smu_ repressor. We also provide evidence of broader effects on *S. mutans* physiology, including slowed growth and disrupted cell morphology when TnSmu1 is activated. In summary, we have discovered an important regulatory module controlling TnSmu1 activation, which extends prior findings made in *ICEBs1*, and illustrates the complex relationship between mobile elements and their hosts.

## RESULTS

### Evidence that *immR*_Smu_ is not an antitoxin

Following our initial observations, we began by investigating the possibility that *immRA*_Smu_ is a toxin-antitoxin module. We reasoned that we would be able to mutagenize both *immR*_Smu_ and *immA*_Smu_ at the same time, as this would not lead to the accumulation of the putative toxin. For other toxin-antitoxin modules (e.g. *mazEF* or *relBE*) double knockout of both genes is possible, whereas single knockout of the antitoxin module is not permitted. Under the conditions tested, we have not been able to obtain a double mutant of *immRA*_Smu_ (Figure S1). However, deletion of the putative metalloprotease, *immA*_Smu_, is permitted by *S. mutans* cells (Figure S1). A lack of viability for a double knockout provides some evidence that *immRA*_Smu_ is not a toxin-antitoxin module. To confirm, we also created over-expression strains in *E. coli* to test the toxicity of each protein. The *immR*_Smu_ and *immA*_Smu_ genes were cloned into a pBAD protein expression vector, where the genes are induced by the addition of the monosaccharide arabinose. When the expression of *immA*_Smu_ was induced, there was only a minor impact on the growth rate of *E. coli*, suggesting that *immA*_Smu_ accumulation in *E. coli* is not toxic (Figure S2). Over-expression of the *immR*_Smu_ protein did cause a moderate growth phenotype in *E. coli*. The growth phenotype was absent when *immR*_Smu_ expression was not induced by arabinose. To summarize, the data do not support the hypothesis that ImmRA_Smu_ constitute a toxin-antitoxin module.

### Phenotypic defects caused by repression of *immR*_Smu_

Next, we wanted to employ CRISPRi as a tool to characterize any phenotypic impacts caused by repression of *immR*_Smu_. Phenotyping is performed by targeting the *immR*_Smu_ gene with a short guide RNA (sgRNA) which acts together with dead Cas9 (dCas9) to block gene transcription; this system is also inducible with xylose [5]. Depletion of *immR*_Smu_ causes a substantial growth defect compared with the control strain (targeting a non-essential gene *lacG*) (Figure 1A). Having shown that *immR*_Smu_ repression leads to a strong growth defect, we next investigated if repression causes any cell morphology impacts. Using transmission electron microscopy (TEM) we were able to gather ultrastructural insights into cells experiencing *immR*_Smu_ depletion. As shown in Figure 1B, when *immR*_Smu_ was depleted, there was a sub-population of sgRNA-*immR*_Smu_ cells with extreme cell morphology defects. Compared to the control strain (sgRNA-lacG) *immR*_Smu_ depleted strains were bloated and appeared to be not dividing or not dividing correctly (at the cell septum) (Figure 1B). Overall, repression of *immR*_Smu_ has a substantial impact on the normal physiology of *S. mutans*.

**Figure 1.**
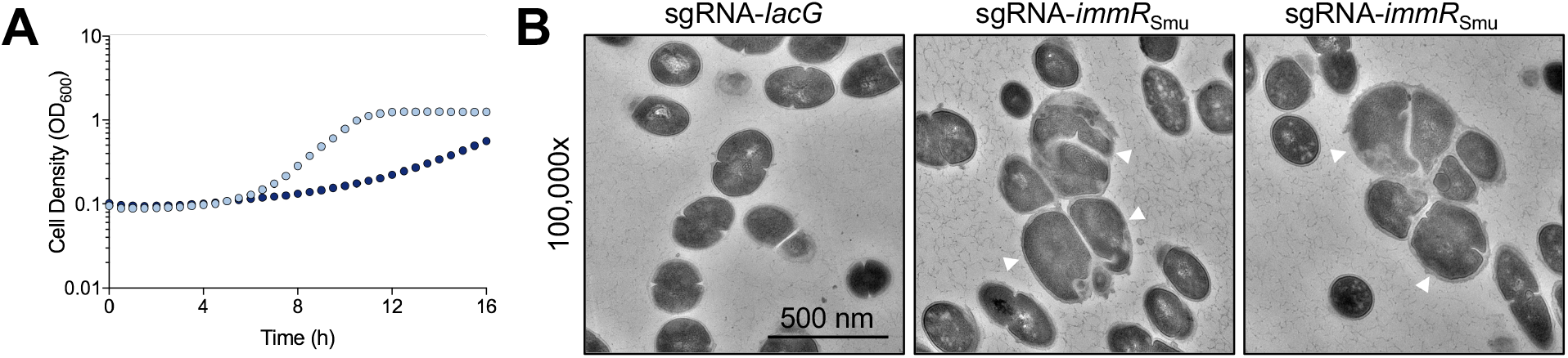
Growth profile and morphological changes in *immR*_Smu_ depleted strains. A CRISPRi strain with an sgRNA targeting *immR*_Smu_ was generated and cultured in CDM containing xylose to repress *immR*_Smu_ gene expression. In these experiments a strain with an sgRNA targeting *lacG* was constructed and serves as a control. (A) Cell densities were measured every 30 min for 16 h with the control strain depicted as light blue dots and the sgRNA-*immR*_Smu_ strain depicted as dark blue dots. (2) Morphological changes were examined with transmission electron microscopy. Depletion of *immR*_Smu_ led to a sub-population of cells to become noticeably larger (white arrows) with aberrant cell division.

### Repression of *immR*_Smu_ with CRISPR inference strongly up-regulates TnSmu1

To begin to understand the function of *immR*_Smu_ we performed RNA-seq profiling of an *S. mutans* strain experiencing knockdown of the *immR*_Smu_ gene using CRISPRi. As shown, when *immR*_Smu_ is repressed by CRISPRi there is a strong growth defect (Figure 1A). RNA was collected from uninduced and induced (with 0.1% xylose) cultures grown to OD_600_ ~0.4, and this RNA was then processed for RNA-seq. The RNA-seq analysis identified 134 genes whose log_2_ fold change was ≥2 after repression of SMU_218 (FDR < 0.05; Table S1). Of these genes, 34 were repressed and 100 were induced (Figure 2A).

**Figure 2.**
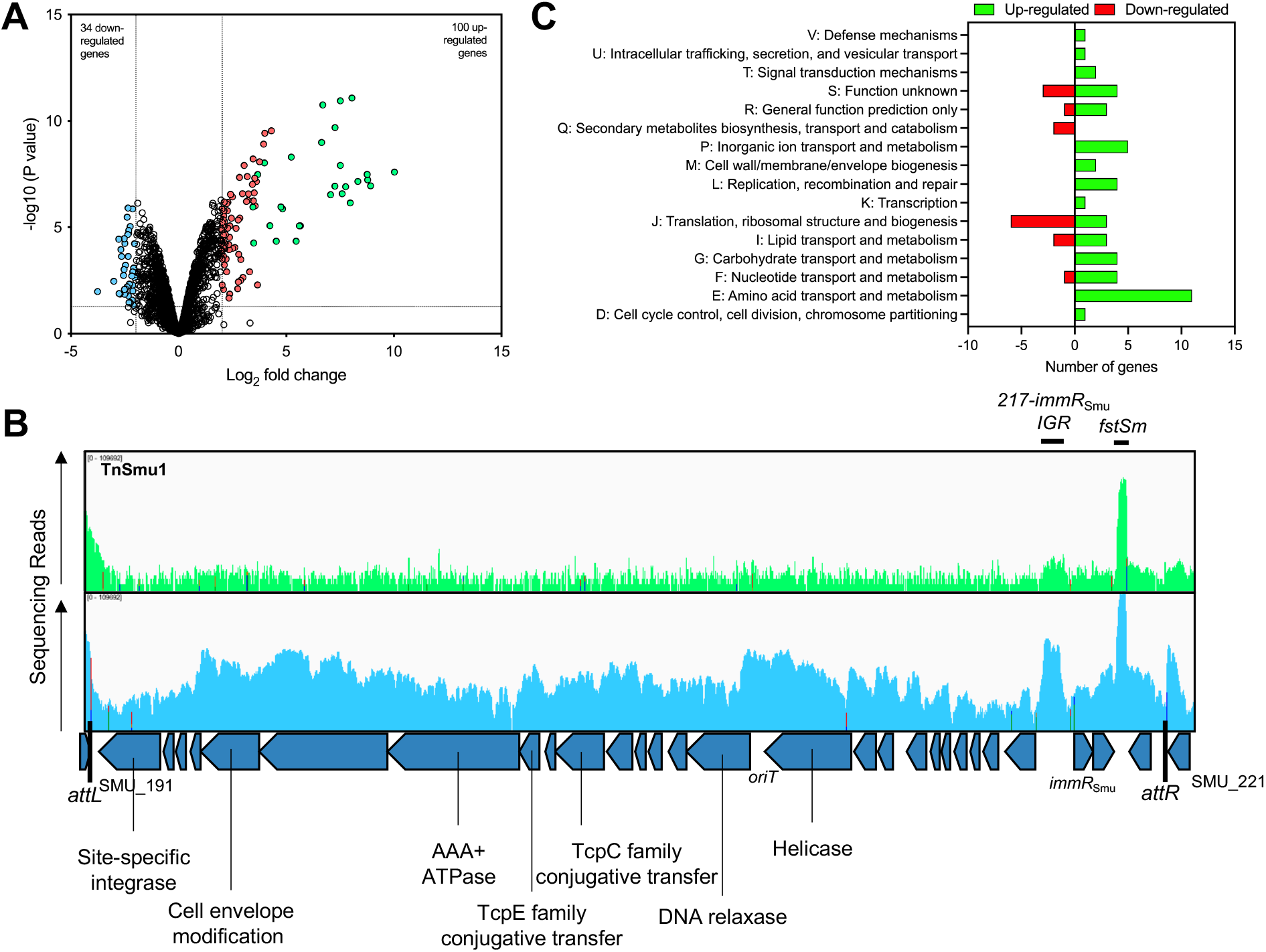
Measuring the *S. mutans* transcriptome response to *immR*_Smu_ repression with CRISPRi-RNAseq. (A) Volcano plot of differential gene expression of *S. mutans* in a CRISPRi strain targeting *immR*_Smu_ (with and without xylose *dcas9* induction). The expression difference was considered significant for a log2 fold change > 2 (vertical dotted line) and p value < 0.05 (horizontal dotted line). TnSmu1 genes are shown as light green dots. (B) RNAseq reads were visualized using the Integrated Genome Viewer, with the control sample shown in green, and the *immR*_Smu_-targeting sample shown in blue. Read coverage is shown across the TnSmu1 mobile element. The *fstSm* and 217-218 intergenic region (IGR) small RNAs are highlighted. (C) The number of genes up- or downregulated with a Cluster of Orthologous Group (COG) assignment is indicated in a bar chart.

There are several notable trends identified by transcriptome analysis of the *immR*_Smu_ depleted strain. Firstly, every gene within the TnSmu1 element is strongly induced (Figure 2A; Table S1). Most TnSmu1 genes were up-regulated by greater than 20-fold, with SMU_204c (hypothetical protein) induced by over 1000-fold (log_2_ fold change = 10.03). Of the twenty most strongly induced genes, all belonged to the TnSmu1 region. Visual inspection of the TnSmu1 region, with reads shown and comparable to a non-induced strain, highlights the strong induction across the element (Figure 2B). This region is predicted to be an ICE and exists in the ICE database ICEberg 2.0 [15] but to our knowledge has not been experimentally verified. To be considered an ICE, these mobile elements must contain several important features: a recombinase (also known as an integrase), a specific site of attachment (typically a tRNA gene), conjugation machinery, a relaxase, and an *oriT* site [16]. Several of these features exist within the TnSmu1 region (Figure 2B). SMU_191c encodes for a putative tyrosine recombinase that may function to catalyze DNA breakage and rejoining. Immediately adjacent to the 5’ end of SMU_191c is tRNA-leu, which is likely the integration site. The attachment (*att*) sites *attL* and *attR* have the sequence CTATACCGGCGGCCG. A putative MOB_T_ family relaxase is encoded by the gene SMU_207c and may function to convert a circular form of TnSmu1 into ssDNA prior to conjugation. Several genes within TnSmu1 are predicted to be members of a type IV secretion system (SMU_196c, SMU_197c, SMU_198c, SMU_199c, SMU_201c, and SMU_208c). A putative *oriT* site is located between SMU_207c and SMU_208c, with the sequence ACCCCCCTATTAGTATCGGGGGG. This shares similarities with the *oriT* site in the ICEs *ICEBs1* (ACCCCCCCACGCTAACAGGGGGGT) and ICE*St3* (ACCCCCGATTTCTAATAGGGGGGT). The relaxase encoded by SMU_207c may recognize this *oriT* sequence, where it would nick the ICE DNA to form transfer DNA (T-DNA). Altogether, TnSmu1 contains many of the genes that are required for an ICE, and may allow excision, replication, transfer, and integration of the element. In addition, strong up-regulation of these genes upon *immR*_Smu_ depletion is suggestive of ImmR_Smu_ acting as a repressor of TnSmu1.

Worth highlighting is that two intergenic regions within TnSmu1 are strongly induced (Figure 2B). The first region is between SMU_217c and *immR*_Smu_, and the second region is between *immA*_Smu_ and SMU_220c. For the first region, we do not currently know what is being produced (*e.g*. small RNA, small peptide, antitermination system *etc*). The second region contains a chromosomally encoded type I toxin-antitoxin system [17]. Expression of a toxic peptide known as Fst-Sm, encoded by this system, is toxic when over-expressed in *S. mutans* (in the absence of the anti-toxin) [17]. The mechanism of action of the toxic peptide is unknown but plasmid encoded Fst peptides affect the integrity of the membrane, cause defects in cell division, and sensitize cells to nisin (by altering the cell membrane) [18, 19].

Several other genes are up or downregulated after *immR*_Smu_ depletion. A broad overview of the effect on the *S. mutans* transcriptome is shown in Figure 2C with categorization by Cluster of Orthologous Groups (COGs). Several genes involved in amino acid transport or metabolism are up-regulated including a cysteine transport operon *tcyDEFGH, argD, cysK, opcD, pepQ, ilvE, gatA, hipO*, SMU_1216c (putative cystine transporter), SMU_1486c (putative histidinolphosphatase), and SMU_1938c (putative methionine transporter). Other transport systems that are also up regulated include *treB* (trehalose PTS), *fruA* (fructose PTS), ammonium transporter (*nrgA*), bacteriocin/competence transporters (*comB*, SMU_1889c, SMU_1897, *comG, mutE2* and *mutG*), *tauC* and *msmK*. The expression of several purine metabolism genes is altered with *purDEK* up-regulated and *purC* down-regulated. All the genes in the Type I-C CRISPR-Cas system are up-regulated when *immR*_Smu_ is repressed. Several genes are downregulated after CRISPRi repression of *immR*_Smu_, many of which are hypothetical proteins. Six translation genes are down-regulated, predominately 50S and 30S ribosomal proteins. Some, but not all, genes located in the genomic island TnSmu2 are downregulated (*mubD, mubC, mubH, mubG*, and *mubE*). This region encodes for a hybrid nonribosomal peptide (NRP) and polyketide (PK) system that produces mutanobactin [20]. A putative acyl carrier protein (SMU_27, *acpP)* is also down-regulated that may participate in PK synthesis [21].

### A bypass of essentiality screen reveals genes that when inactivated allow the deletion of *immR*_Smu_

Under the conditions we have tested, we are not able to obtain mutants of *immR*_Smu_ or *immRA*_Smu_ but deletion of *immA*_Smu_ is permitted (Figure S1). We hypothesized that deletion of the *immRA*_Smu_ module might be possible if introduced into a transposon library previously generated by our laboratory [4]. This is because the Tn-seq library might contain mutants that interrupt a gene responsible for *immRA*_Smu_ essentiality and thus would tolerate *immRA*_Smu_ deletion. This is known as bypass of essentiality (BoE), with systematic studies of essential gene bypass having been conducted in yeast [22, 23]. BoE has been shown for several essential genes in bacteria including *zipA*, c-di-AMP null mutants, RNase E, *bamD*, and *ftsH* [24–28]. Utilizing a BoE screen, where we transformed an *immRA*_Smu_ deletion construct into the Tn-seq library, we were able to isolate eight double mutants capable of tolerating *immRA*_Smu_ deletion (Figure 3A). These mutants were then subjected to complete genome sequencing, to identify the location of the transposon insertion and identify suppressor mutations in the event these were also generated. Genome sequencing revealed the transposon insertions to be within the following regions: SMU_197c, SMU_201c (isolated twice), SMU_141c, *recN, perR*, the *murE* and SMU_1678 intergenic region, and the intergenic region between 16S rDNA and SMU_1750c (Table S2). The growth phenotypes of these strains were variable (Figure 2B). The intergenic transposon insertion into *murE* caused a considerable growth phenotype, as did an insertion in SMU_197c. Certain double mutant isolates grew almost the same as wild-type *S. mutans*, including *peR*::Tn, *recN::Tn* and SMU_141c::Tn.

**Figure 3.**
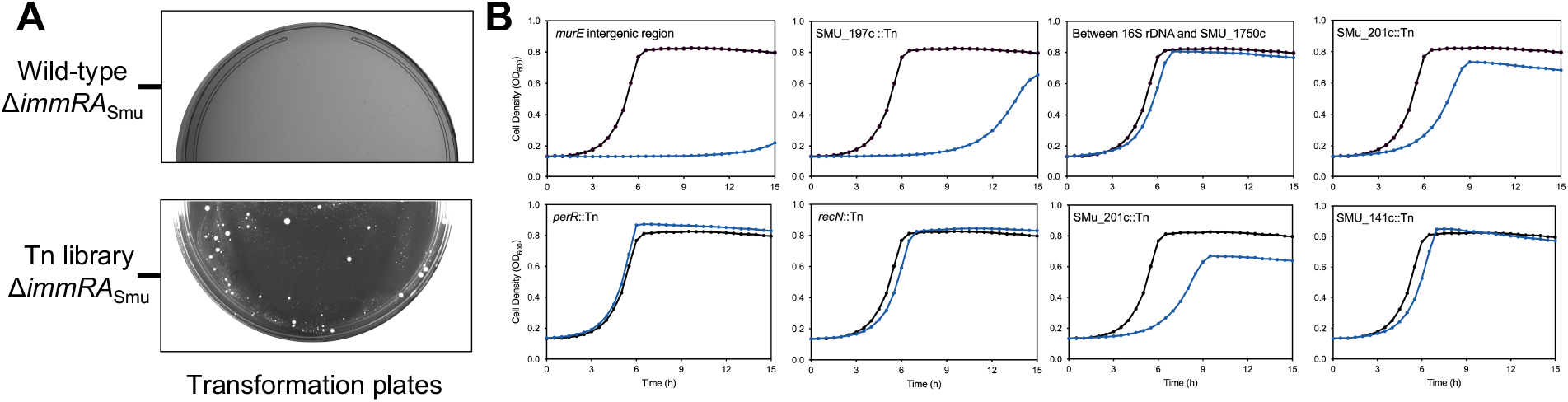
A transposon interaction screen identifies gene mutations that bypass the essentiality of *immR*_Smu_. (A) An *immRA*_Smu_::*aphA3* mutagenesis fragment was transformed into a *S. mutans* Tn-seq library with selection on antibiotic-containing agar. Deletion of *immRA*_Smu_ is not possible in a wild-type strain of *S. mutans* (top panel). White *S. mutans* colonies were obtained when *immRA*_Smu_ was transformed into the Tn-seq library (bottom panel). (B) After genome sequencing analysis, several strains were identified that allow *immRA*_Smu_ deletion. Each strain, with the corresponding transposon insertion site, is shown in a single growth analysis chart (dark line, wild-type strain; light blue line, double mutant strain).

We hypothesized that because of the strong selective pressure placed on cells during transformation, mutants could be created that are experiencing a duplication event; this is the loss of the essential module *immRA*_Smu_ but a copy of *immRA*_Smu_ is still present. To account for this, we used PCR to visualize the size of the *immRA*_Smu_ region. Gene duplication events occurred in the strains *perR*::Tn, *recN*::Tn, and the strain with a transposon insertion between 16S rDNA and SMU_1750c (Figure S3; Table S2). Double mutants with an *immRA*_Smu_ gene duplication grew better than strains that fully lacked *immRA*_Smu_ (Figure 2B). For the other five mutants, only one band for *immRA*_Smu_ was amplified and the size was as expected for a deletion mutant. As an additional control, PCR was also conducted to confirm that the Magellan6 transposon was inserted in the regions identified by whole genome sequencing. Transposon insertions were apparent for all strains except for SMU_141c::Tn where the results were inconclusive (Figure S4; Table S2).

### Evidence of a circular intermediate form of TnSmu1

Next, we noticed that in several double mutant backgrounds, read coverage (after genomic sequencing) was significantly elevated for a region between SMU_191c and SMU_220c (Figure 4; Figure S5). Read coverage depth in this region was ~300 for *S. mutans* UA159 but was elevated to as much as ~5000 reads in double mutant backgrounds. This increased read depth corresponds exactly to a region of the genome that is predicted to contain the putative ICE, TnSmu1. When excised from the host chromosome ICEs exist as circular DNA molecules [29]. For some ICEs these circular intermediate forms are capable of rolling-circle replication [30]. Increased read coverage depth suggests that loss of *immR*_Smu_ is leading to excision and circularization of TnSmu1. In addition to the observation that sequencing reads increase when *immR*_Smu_ is deleted, there was also evidence of unexpected genomic junctions in strains of *S. mutans* missing the *immR*_Smu_ gene. There was evidence from the genome sequencing that *attL* and *attR* are combining, which is leading to a junction forming between SMU_191c and SMU_220c. This is additional evidence that TnSmu1 is excising from the host chromosome and forming a circular DNA molecule. To confirm these observations, we designed primers to test excision and circularization. For excision, these primers will only amplify a PCR product if TnSmu1 is no longer present on the host chromosome. For circularization, a PCR product will form if there is a junction between the ends of the ICE (*i.e*. SMU_191c and SMU_220c). Excision and circularization of TnSmu1 in wild-type *S. mutans*, grown to mid-log, occurs close to the limit of detection for the assay (Figure 5A and B). However, excision and circularization of TnSmu1 is apparent in *immR*_Smu_ double mutant strains. For Δ*immR*_Smu_ strains carrying transposon insertions within putative TnSmu1 conjugation genes (SMU_197c and SMU_201c) there are multiple copies of TnSmu1 per cell. In these strains excision is occurring 1000-fold more compared to wild-type *S. mutans*. We also observed circularization/excision in two strains experiencing duplication of *immR*_Smu_, *perR*::*Tn* and *recN*::*Tn*. We chose to examine these strains because they are much healthier than the strains with disrupted conjugation genes (Figure 3B). For these two strains, circularization of TnSmu1 was occurring but excision of the element was measured close to the limit of detection for the assay. Taken together, our observations of excision and circularization are consistent with TnSmu1 having this important life-cycle feature of ICEs.

**Figure 4.**
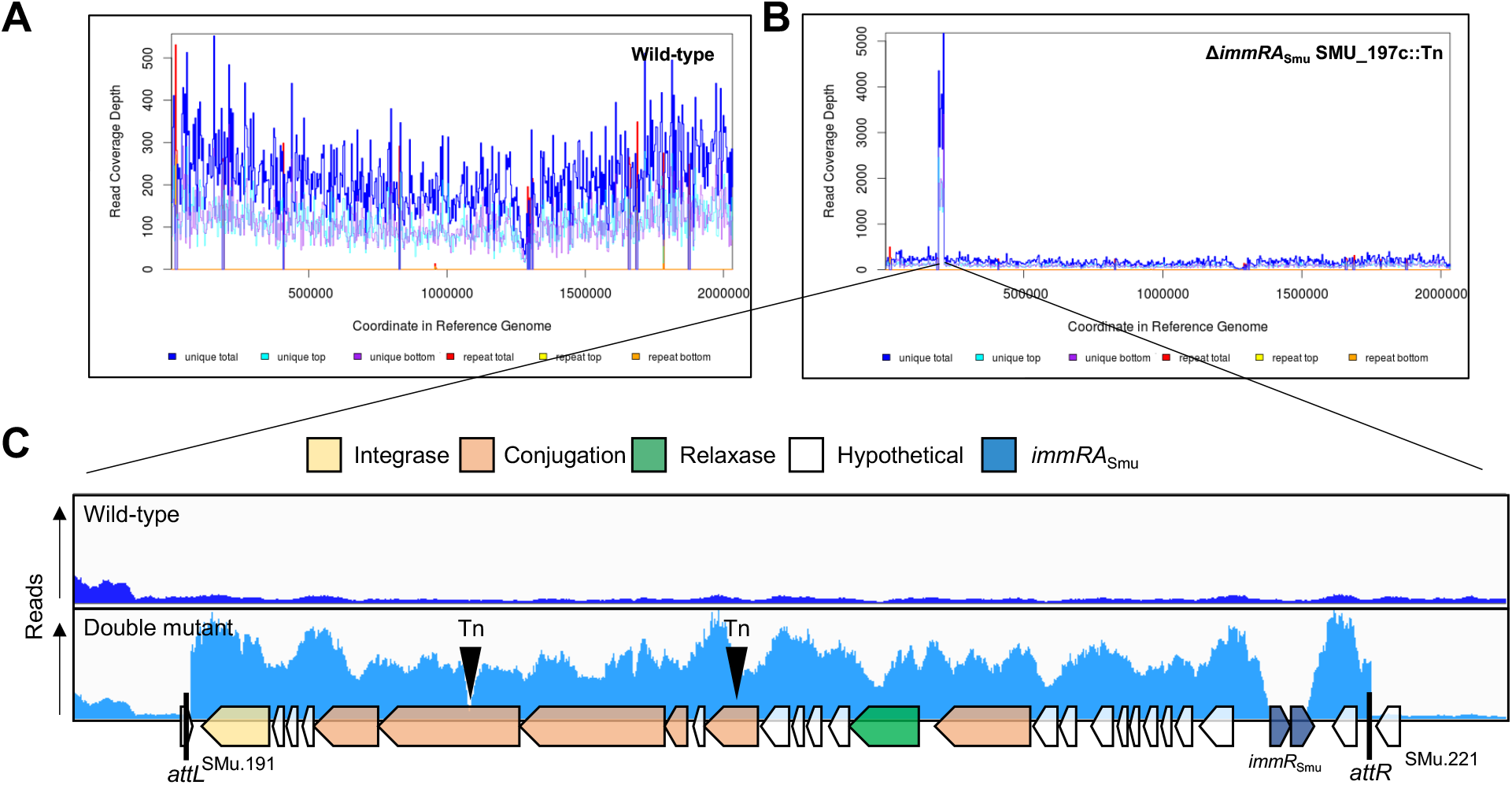
Mutations that bypass the essentiality of *immRA*_Smu_ cause an increase in genome sequencing reads from the ICE TnSmu1. Read coverage depth for both a wild-type (A) and a double mutant strain Δ*immRA*_Smu_::*aphA3* SMU_197c::Tn (B) is shown. Genome sequencing read depth was substantially higher across the TnSmu1 region in the double mutant background. (C) Sequencing reads were visualized with the Integrated Genome Viewer for only the TnSmu1 region (and immediate up- and downstream genes). Read coverage is ca. 10-fold higher across the entire TnSmu1 region, except for the deleted *immRA*_Smu_ genes. The location of each transposon insertion, within putative conjugation genes, which allowed *immRA*_Smu_ deletion is shown as upside-down triangles. The TnSmu1 region is annotated with putative functions related to excision/recombination (integrase), conjugation, T-DNA formation (relaxase), strand nicking (*oriT*) and attachment sites (*attL* and *attR*).

**Figure 5.**
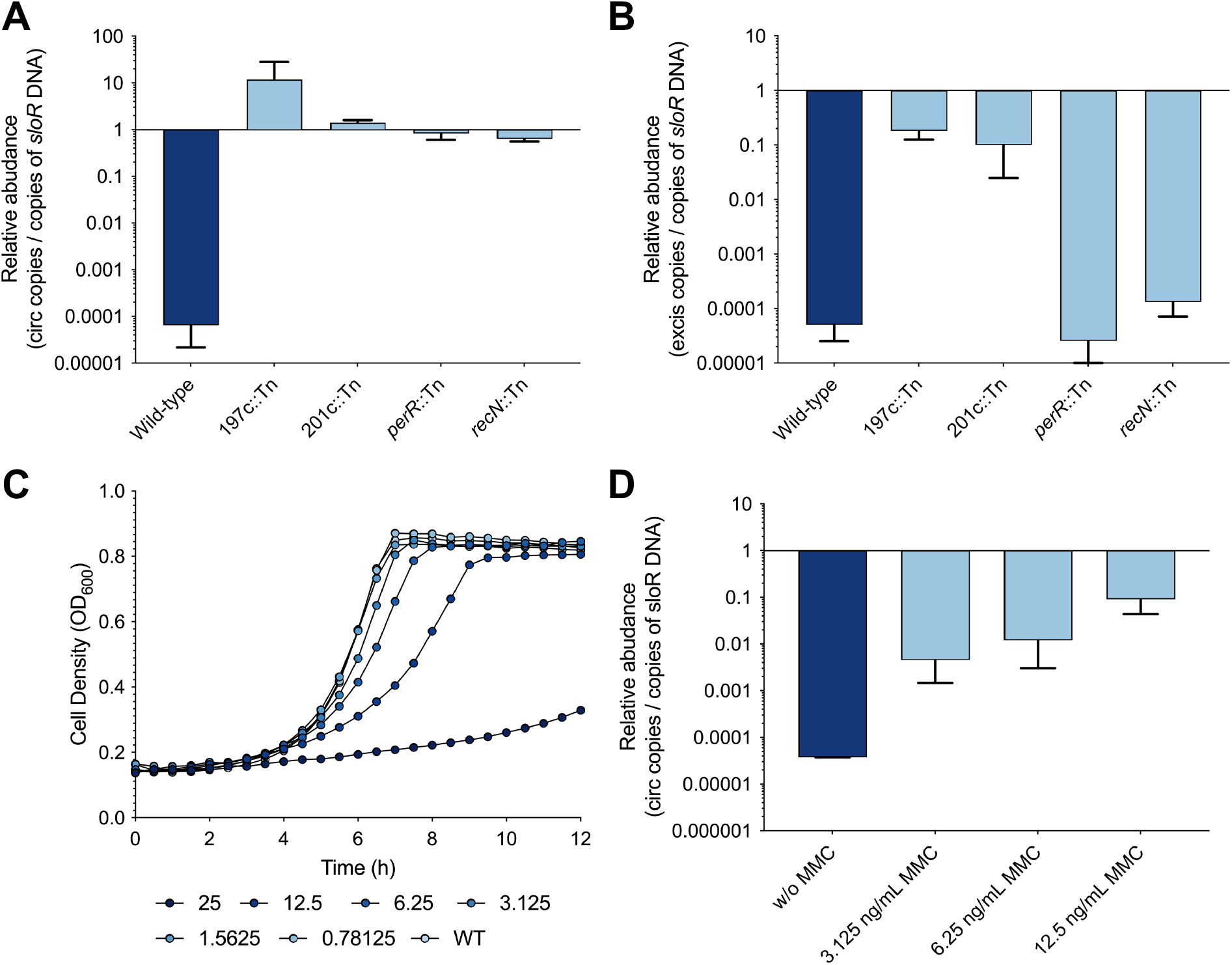
TnSmu1 is capable of circularization and excision. Quantitative PCR was employed to measure the relative abundance of circular (A) and excised (B) copies of TnSmu1. The number of copies was compared to a nearby single-copy chromosomal gene, *sloR*. Induction of TnSmu1 circularization was also measured by inducing TnSmu1 with the genotoxic agent mitomycin C (MMC). The impact of different concentrations of MMC was quantified using a growth assay (C), and then different doses of MMC were used to induce TnSmu1 circularization (D).

### Induction of TnSmu1 by the genotoxic agent mitomycin C

The *immRA*_Smu_ module shares similarities to cI-like phage repressors that are sensitive to DNA damage (via induction of the SOS response) [31]. Other ICEs contain similar regulatory modules, *ICEBs1* being the most extensively characterized but also ICE*St3* and SXT [10, 32, 33]. To determine if DNA damage was an inducer of TnSmu1, possibly via cleavage of ImmR_Smu_, we incubated *S. mutans* UA159 in the presence of genotoxic agent mitomycin C (MMC). When treated with MMC there was a clear dose-dependent response, with clear growth hindrance at 25 ng/mL (Figure 5C). Induction of TnSmu1 circularization was investigated at MMC concentrations of 3.125 ng/mL, 6.25 ng/mL and 12.5 ng/mL. With increasing concentrations of MMC, the relative abundance of circular TnSmu1 increased in a dosedependent manner (Figure 5D). At a concentration of 12.5 ng/mL not all cells have a circular copy of TnSmu1 but circularization is 1000-fold above levels seen in untreated *S. mutans* cells.

### Transcriptome analysis in *S. mutans* strains lacking *immR*_Smu_

In order to provide further evidence that *immR*_Smu_ represses activation of TnSmu1, we examined the transcriptomes of two strains generated during the BoE screens. RNA was extracted from SMU_197c::Tn Δ*immRA*_Smu_ and SMU_201c::Tn Δ*immRA*_Smu_ cultures grown to OD_600_ ~0.4, and this RNA was then processed for RNA-seq. For the SMU_197c::Tn Δ*immRA*_Smu_ strain RNA-seq analysis identified 54 genes whose log_2_ fold change was ≥2 (FDR < 0.05; Table S3). Of these genes, 7 were repressed and 47 were induced (Figure 6A). Similar results were obtained for the SMU_201c::Tn Δ*immRA*_Smu_ strain with RNA-seq analysis identifying 51 genes whose log_2_ fold change was ≥2 (FDR < 0.05; Table S4); 7 genes were repressed and 44 were induced (Figure 6A). As with CRISPRi repression of *immR*_Smu_, TnSmu1 expression is significantly induced in both Δ*immR*_Smu_ strains. In both strains SMU_205c (hypothetical protein residing in TnSmu1) was the most strongly up-regulated gene by 968-fold (SMU_201c::Tn Δ*immR*_Smu_) and 399-fold (SMU_197c::Tn Δ*immR*_Smu_). Next, we generated a heat map of selected genes with the aim of comparing gene expression between Δ*immR*_Smu_ strains and the sgRNA-218 strain (Figure 6B). Notably, up-regulation of TnSmu1, the Type I-C CRISPR-Cas system and the CslAB transporter system is consistent among all three strains. In addition, SMU_40 (hypothetical protein with a RelE toxin-antitoxin system domain) is moderately up-regulated (~5-fold) across the three strains. For the malolactic fermentation (*mle*) locus (SMU_137 to SMU_141)[34], downregulation (~10-fold) was observed in the Δ*immR*_Smu_ strains, and up-regulation (~3.5-fold) in the sgRNA-218 strain. The *mle* locus contains a malolactic enzyme (*mleS*; SMU_137), a malate permease (*mleP*; SMU_138), an oxalate decarboxylase (*oxdC*; SMU_139), a glutathione reductase (*gshR;* SMU_140) and a hypothetical protein (SMU_141); the role of this operon is in malolactic fermentation (conversion of L-malate to L-lactate). Taken together, this additional RNA-seq analysis shows that loss of Δ*immR*_Smu_ is leading to activation of TnSmu1. In addition, there is a consistent trend showing activation of the Type I-C CRISPR-Cas system and the *cslAB* transporter system.

**Figure 6.**
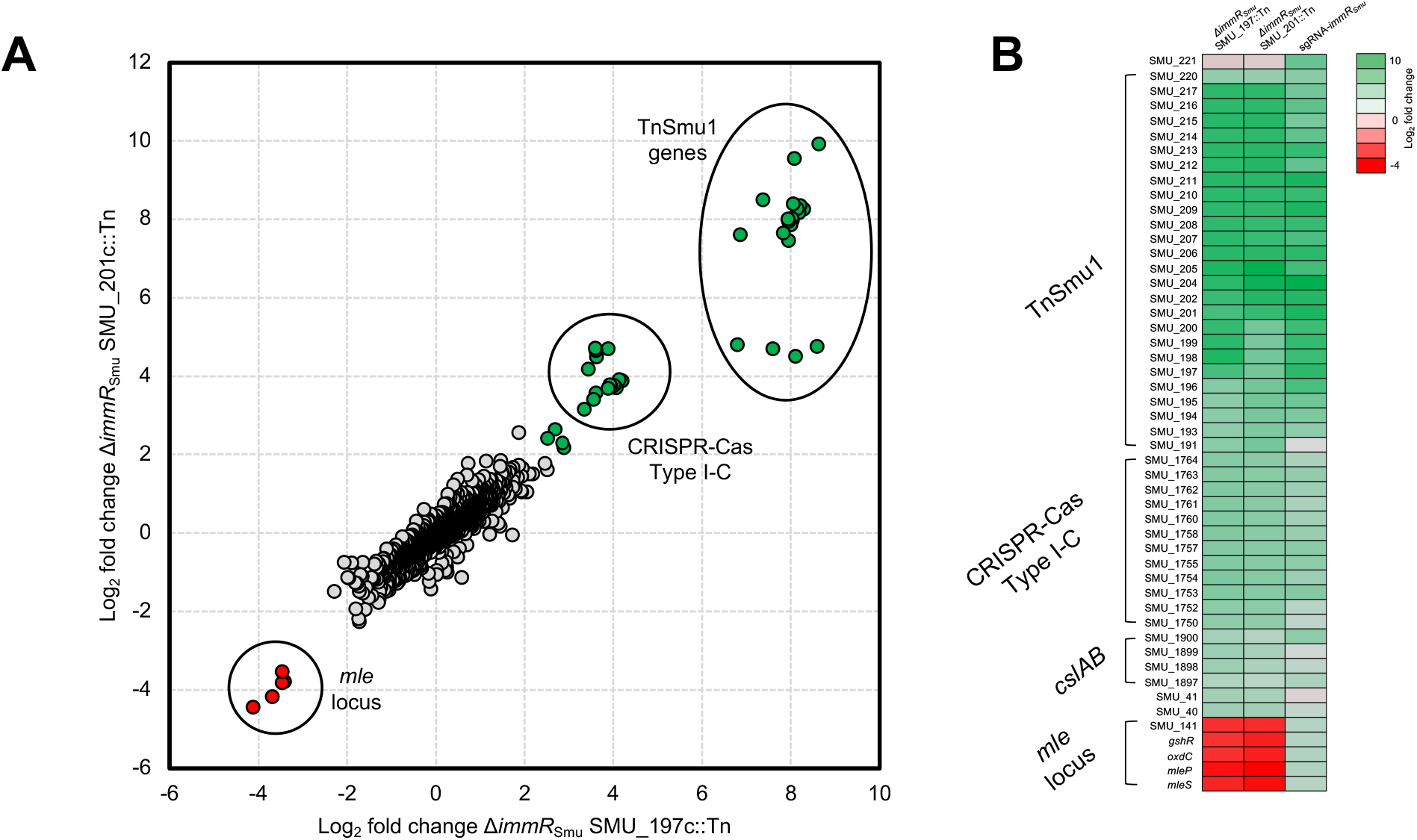
RNA-seq analysis of strains lacking *immRA*_Smu_. RNA was extracted from SMU_197c::Tn Δ*immRA*_Smu_ and SMU_201c::Tn Δ*immRA*_Smu_ cultures grown to OD_600_ ~0.4, and this RNA was then processed for RNA-seq. (A) Comparison of gene expression (log2 fold change) between SMU_197c::Tn Δ*immRA*_Smu_ and SMU_201c::Tn Δ*immRA*_Smu_. Pearson’s correlation coefficients between transcriptomes are presented. Gene expression of TnSmu1, Type I-C CRISPR-Cas and the *mle* locus are highlighted on the chart. (B) Heat map of selected bacterial transcripts that are enriched or depleted in the chosen strains. Colors represent the log2 fold change compared to control conditions with a key shown to the right of the heat map.

### Distribution and conjugative transfer of TnSmu1 genes between *S. mutans* strains

To begin to understand if TnSmu1 is capable of conjugative transfer, we wanted to determine if closely and distantly related strains of *S. mutans* carry this ICE. Notably, we have recently discovered that *S. mutans* strains carry CRISPR spacers against this element [35]. This finding suggests that *S. mutans* attempts to prevent transfer/acquisition of TnSmu1 via CRISPR-Cas mediated immunity, and it also provides evidence of potential transfer between strains. Figure 7 shows strains that carry TnSmu1 genes and is organized by strain relatedness. Distantly related strains, such as smu342 and UA159, carry full length TnSmu1 elements, which might be indicative of conjugal transfer. Although many *S. mutans* strains carry TnSmu1 genes, only nine strains carry the full element as organized on *S. mutans* UA159. ICEs have plasticity and lose/gain genes, often interacting with other mobile elements such as transposons, lysogenic phages and genomic islands. It is therefore not surprising that the structure of TnSmu1 changes as it transfers within the *S. mutans* species, particularly if its expression is lethal in other *S. mutans* hosts. Under normal conditions we anticipate that a very small sub-fraction of *S. mutans* cells would express TnSmu1 because of ImmR_Smu_-mediated repression of the element. It is therefore likely that transfer of TnSmu1 from a donor strain to a recipient strain occurs at very low efficiency. Similar ICEs in other Firmicutes bacteria, like *ICEBs1* and Tn*916*, exhibit considerable differences in conjugation rates. In the future we plan to explore conjugative transfer of this element between strains in greater detail.

**Figure 7.**
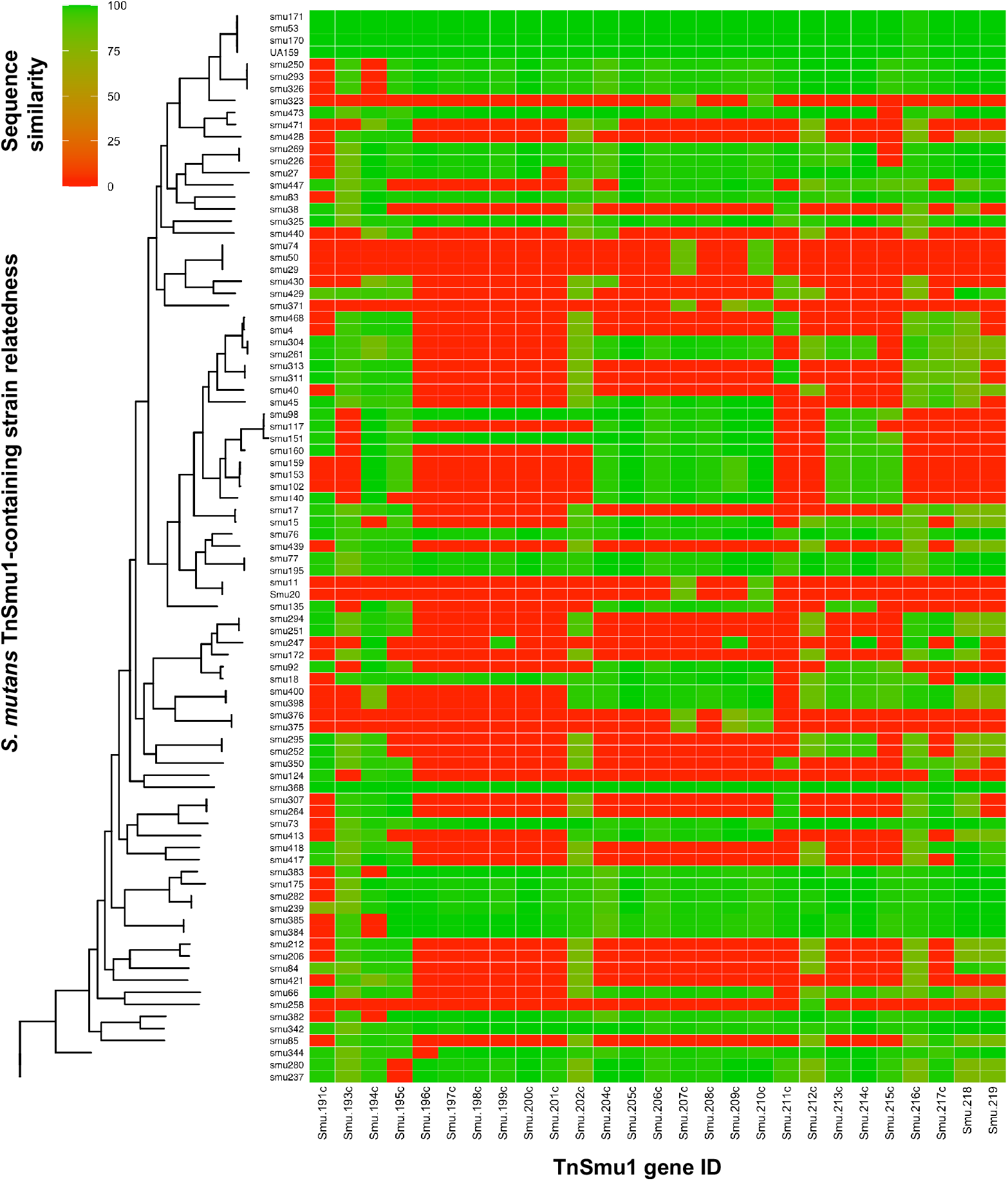
TnSmu1 gene distribution across sequenced *S. mutans* genomes. *S. mutans* strains were searched for TnSmu1 genes, comparing to those carried on *S. mutans* UA159. Strains are organized by strain relatedness (*y*-axis) and gene ID is shown on the *x*-axis (SMU_191c to SMU_219). The percent sequence similarity for each gene is shown (compared to the gene on *S. mutans* UA159 TnSmu1). Red areas depict a lack of a TnSmu1 gene on a strain genome.

## DISCUSSION

In this study, we sought to determine the role of SMU_218/*immR*_Smu_, a gene we previously identified as being ‘essential’ or non-mutable by Tn-seq. We hypothesized based on genomic context and protein domain similarities that *immR*_Smu_ was either an antitoxin gene in a toxin-antitoxin system, or a transcriptional regulator involved in activation of the ICE TnSmu1. We provide evidence that *immR*_Smu_ is not an antitoxin. Instead, we have concluded that *immR*_Smu_ is a repressor that keeps TnSmu1 silent, except under certain environmental conditions. In exploring *immR*_Smu_ we provide evidence that TnSmu1 can be induced, that it produces a circular intermediate, and the element, or parts of it, are distributed among *S. mutans* strains. We also uncovered that activation of TnSmu1 has a considerable impact on *S. mutans* physiology, leading to changes in gene expression, slowed growth and abnormal cell morphology.

Our conclusion that *immR*_Smu_ is a repressor involved in the activation of TnSmu1 is based on several lines of evidence. Firstly, BLAST and other related bioinformatics searches show that *immR*_Smu_ is a predicted helix-turn-helix DNA binding protein, and the downstream gene *immA*_Smu_ is a predicted Zn^2+^ metalloprotease. BLAST searches also reveal that *immRA*_Smu_ share similarities with other described transcriptional regulator and metalloprotease regulatory modules. Most importantly, the mobile element *ICEBs1* contains a repressor, ImmR, that is cleaved by a protease ImmA, in a two gene organization like *immRA*_Smu_ but in a different location within the ICE (Figure 8A) [11, 33]. Due to the similarities of the system described here, and ICE*Bs1 immRA*, we have named SMU_218 *immR*_Smu_ and SMU_219 *immA*_Smu_. Activation of ICE*Bs1* occurs when ImmR is cleaved by the protease ImmA, either through a cell-cell signaling pathway involving RapI and PhrI or via a RecA-dependent DNA damage response [11, 33]. TnSmu1 does not appear to contain a RapI/PhrI cell-cell signaling system but additional regulatory systems on top of *immRA*_Smu_ cannot be ruled out, as TnSmu1 contains several hypothetical genes with unknown functions. ImmRA-like systems are encoded on other putative and studied mobile genetic elements. ICE*St1* and ICE*St3* contain a regulation module with a ImmR-like gene (*arp2)*, a metalloprotease gene (*orfQ*) and a cI gene (*arp1*) [14]. In this module the metalloprotease gene, *orfQ*, is not downstream of the *immR*-like gene, *arp2*, but as with ICE*Bs1* these elements are activated by DNA damage, which is probably regulated via repressor proteolysis [14]. An ImmRA-like system is also required for the activation of a staphylococcal pathogenicity island SaPI3 [36]. For this regulation module, ImmR repression is partially alleviated when Sis (<u>S</u>aPI <u>i</u>nducer of <u>S</u>aPIs), a protein produced by other SaPIs, binds to ImmR but full activation only occurs with ImmA-mediated proteolytic cleavage of ImmR [36]. ImmRA modules are also encoded by lysogenic phages known to be activated by DNA damage [11]. These modules are therefore a common regulatory system governing activation of mobile genetic elements, and several have additional regulatory complexities in addition to ImmRA.

**Figure 8.**
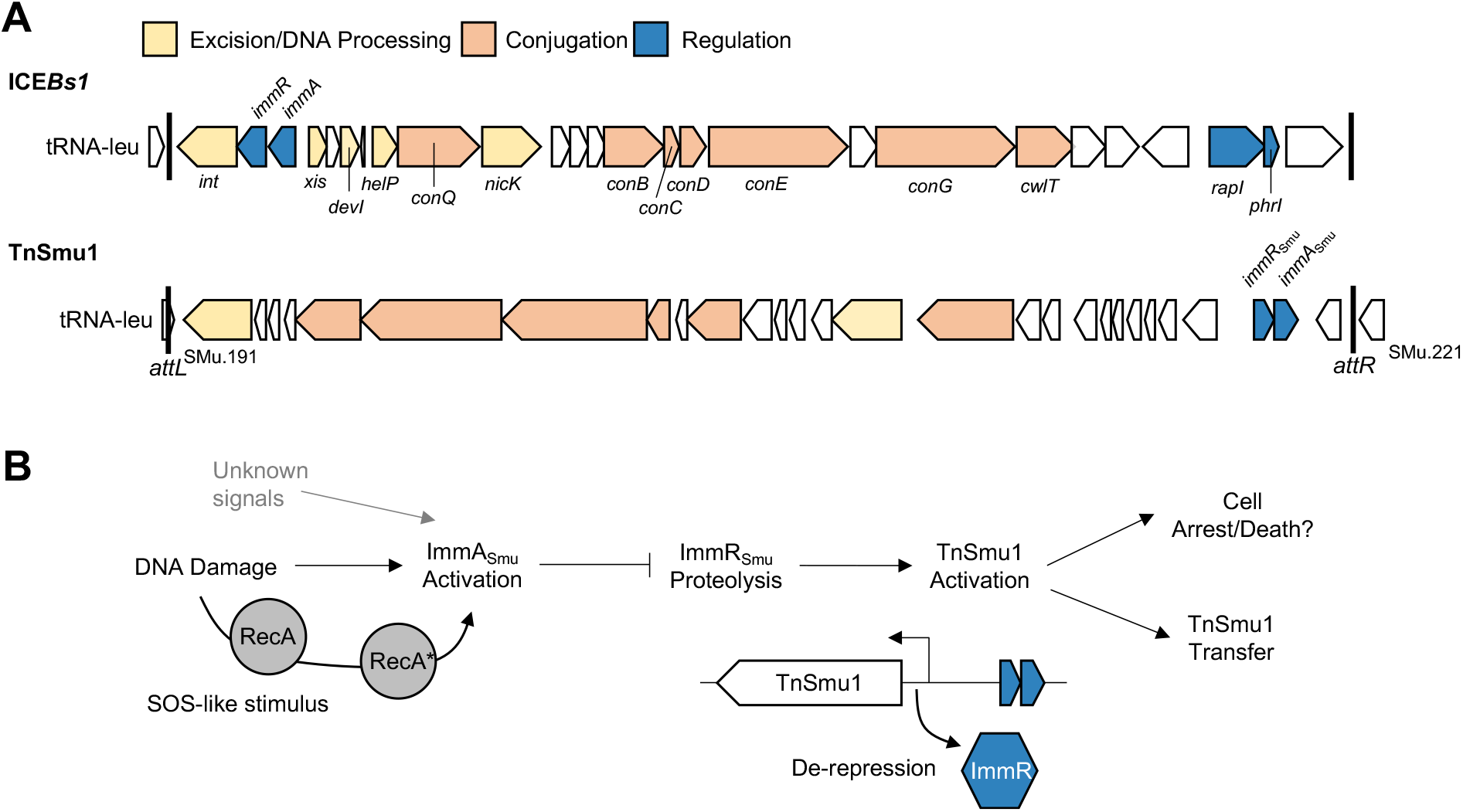
Comparison of *ICEBs1* with TnSmu1 and proposed model for ImmRA_Smu_ regulation of TnSmu1. (A) The genetic organization of *ICEBs1* and TnSmu1 is shown. Genes are colored according to function with their names above or below. Genes with unknown functions are colored white. Both ICEs integrate at tRNA-leu. Note that the *immRA*_Smu_ regulatory module is in a different location compared with *ICEBs1*. (B) A diagram depicting the ImmRA_Smu_ regulatory pathway for TnSmu1 activation is shown. Our model draws heavily from *ICEBs1* ImmRA, and closely related systems, where DNA damage leads to activation of the protease ImmA. Activated ImmA cleaves the ImmR repressor leading to activation of TnSmu1. It is likely that stochasticity causes sub-population responses so that TnSmu1 activation can lead to transfer of the element, cell arrest/cell death and other phenotypes. The SOS pathway, leading to RecA-mediated activation of ImmA_Smu_, is not well known in *S. mutans* and will need to be investigated further. Yet to be discovered signals may also lead to ImmA_Smu_-mediated cleavage of ImmR_Smu_.

Aside from inducing excision of ICE elements, there is evidence that similar transcriptional regulator and metalloprotease modules are encoded in other systems. Recently, a system that parallels ImmRA was identified on CBASS (cyclic oligonucleotide based anti-phage signaling system) anti-phage systems [37]. Called CapH/CapP, this module regulates expression of a CBASS system in response to DNA damage. The transcriptional regulator CapH represses CBASS transcription until it is cleaved by the metallopeptidase CapP, which is stimulated by the presence of single-stranded DNA. Once the CBASS system is derepressed a cell killing pathway is induced which kills the host bacteria. Another ImmRA-like system, known as DdrO/IrrE, is encoded by *Deinococcus* spp. [38]. This system regulates a DNA damage response, with the repressor DdrO being cleaved by IrrE during radiation, leading to expression of DNA repair genes and an apoptotic-like cell death pathway [38]. There is a common theme among the ImmRA, CapHP and DdrO/IrrE modules in that they cause expression of genes upon sensing of DNA damage, with cleavage of a transcriptional repressor by an activated metalloprotease. In addition, both CapHP and DdrO/IrrE lead to host genome killing. That these systems have pathways that lead to cell-killing is notable because activation of TnSmu1 (via CRISPRi) caused growth defects and abnormal cell morphology. However, additional studies are warranted to determine if this is mediated directly or indirectly by TnSmu1.

Additional evidence that *immRA*_Smu_ is a regulatory module was provided by RNA-seq which helped to identify genes activated in response to either depletion or deletion of *immR*_Smu_. When *immR*_Smu_ is repressed or deleted there is a clear and strong up-regulation of TnSmu1-associated genes by as much as 1000-fold. Although derepression of TnSmu1 genes by ImmR_Smu_ would be most strongly confirmed by DNA-binding assays (*e.g*. electrophoretic mobility shift assay), activation of TnSmu1 genes upon loss of the ImmR repressor is consistent with similar systems described in ICE*Bs1*, CapH/CapP and DdrO/IrrE. In addition, deletion of *immR*_Smu_ was found to activate excision/circularization of TnSmu1. Again, this would be consistent with a hypothesis that loss of the repressor leads to constitutive derepression of TnSmu1 followed by activation of the mobile element. Based on ICE*Bs1* ImmRA (and similar systems) we predict that ImmA_Smu_ is a metalloprotease that is activated by DNA damage, leading to cleavage of ImmR_Smu_ and derepession of TnSmu1. Although we do not provide direct evidence confirming this hypothesis, we do show that TnSmu1 circularization is induced in response to DNA damage by the genotoxic agent MMC. Taken together, activation by TnSmu1 as shown in RNA-seq studies and in response to MMC provides reliable evidence that ImmRA_Smu_ functions as a regulatory module which has similarities to ICE*Bs1* ImmRA.

We were initially drawn to studying *immR*_Smu_ because of Tn-seq data showing that the gene was ‘essential’. With the evidence gathered here we are able to conclude that *immR*_Smu_ is not a classical essential gene and does not participate in a core, required for survival, biological pathway. Instead, *immR*_Smu_ appears ‘essential’ because deletion of *immR*_Smu_ activates TnSmu1 leading to excision and eventual loss of the mobile element from the cell; thereby not making it possible to select a deletion strain. The observation that inactivation of nearby conjugation-related genes (SMU_197c and SMU_201c) allows *immR*_Smu_ deletion provides evidence for this hypothesis. Here, disruption of the conjugation genes likely leads to TnSmu1/Δ*immR*_Smu_ becoming ‘trapped’ in the cell, and selectable through antibiotic cassette replacement. However, we were able to describe significant impacts on cell growth, cell morphology and the transcriptome when *immR*_Smu_ was depleted with CRISPRi. Despite in many cases ICEs carrying beneficial genes, such as virulence factors or antibiotic resistance genes, expression of these elements has been found to have a major impact on certain hosts. Induction of ICE*clc* in *Pseudomonas* spp. causes a sub-population of cells to differentiate into “transfer competent” cells that have arrested cell growth and cell lysis [39]. Activation of Tn*916* causes severe cell growth defects in *Enterococcus faecalis* and *Bacillus subtilis* [40]. Tn*916* only activates in a small percentage (0.1-3%) of cells and the lethality of Tn*916* was discovered by activating it in a much larger proportion of cells. The deleterious effects of Tn*916* were less impactful in cells lacking conjugation genes that reside in Tn*916*, and a gene, *yqaR* found within a defective phage-like element *skin* [40]. Intriguingly, Tn*916* can also cause cell death in *E. faecalis* and *B. subtilis*, species that do not encode for a *yqaR*-like gene. Future studies of the growth arrest/cell death mechanism in TnSmu1 would be helpful in understanding this generally conserved process for ICE transfer, and whether it is beneficial or costly for transfer efficiency.

We expect that under most conditions TnSmu1 will be quiescent because of ImmR_Smu_-mediated repression of the mobile genetic element. This silence helps maintain vertical transmission of the element during non-stressful conditions. We anticipate that activation of TnSmu1 occurs when ImmA_Smu_ becomes active and cleaves ImmR_Smu_ leading to de-repression of TnSmu1 (Figure 8B). As with other ICEs, such as ICE*Bs1*, DNA damage causes activation of TnSmu1, as shown here with MMC treatment. For ICE*Bs1*, during DNA damage, ImmA is stimulated via an SOS (save our souls) response that causes RecA to activate ImmA cleavage of ImmR [11, 33]. Although our model of ImmRA_Smu_-mediated regulation of TnSmu1 includes activation of ImmA by RecA, there may be substantial differences in the SOS response between *S. mutans* and *B. subtilis*. *Streptococcus* spp. have generally been thought to lack a classical SOS response because of work performed in *Streptococcus pneumoniae*, which lacks a LexA homologue, where genetic competence plays a central role in responding to DNA damage [41]. However, other streptococci such as *Streptococcus thermophilus* encode LexA-like repressors and competence development is antagonistic to the SOS response [42]. The *S. mutans* UA159 genome encodes two LexA-like repressors, SMU_1398 (*irvR*) and SMU_2027 (*hdiR*). IrvR does not participate in the SOS-response and instead is a stress-responsive biofilm regulator [43]. In comparison, *hdiR* is induced by SOS stress (MMC) in a *recA*-dependent manner [43]. Genetic competence could play a role in an SOS-like response because *recA* is up-regulated as part of the late competence regulon [44]. Notably, other pathways may be involved in TnSmu1 regulation as RNA-seq studies show TnSmu1 up- or down-regulation in different conditions or mutant backgrounds [45–49]. Although our core model of TnSmu1 activation via ImmR_Smu_ proteolysis borrows heavily from the ICE*Bs1* system we anticipate notable differences in how ImmA_Smu_ becomes active.

To conclude, we show that *S. mutans* contains an ICE, TnSmu1, whose activation is likely controlled by an ImmRA-like regulatory module. Supporting this hypothesis is strong up-regulation of TnSmu1 genes and excision/circularization of the element upon loss or depletion of *immR*_Smu_. The mobile element is also inducible via DNA damage, which is a conserved feature of ImmRA-like regulatory systems. Future studies will be directed to better understanding the steps involved in TnSmu1 activation, as well as identifying the mechanism that causes activation to be deleterious to host fitness and cell biology.

## METHODS

### Bacterial strains and culture conditions

*Streptococcus mutans* strains were cultured from single colonies in Brain Heart Infusion (BHI, Difco) broth. Unless otherwise stated *S. mutans* was routinely cultured at 37°C in a 5% CO_2_, microaerophilic atmosphere. *Escherichia coli* strains were routinely cultured in LB broth (Lennox formula; 10 g/L tyrptone, 5 g/L yeast extract and 5 g/L NaCl) at 37°C with aeration. Antibiotics were added to growth media at the following concentrations: kanamycin (1.0 mg/mL for *S. mutans*, 50 μg/mL for *E. coli)*,spectinomycin (1.0 mg/mL for *S. mutans)*, ampicillin (100 μg/mL for *E. coli*). A list of strains and plasmids (Table S5) and oligonucleotide primers (Table S6) can be found in the supplementary material.

### Gene mutagenesis and plasmid cloning

Standard DNA manipulation techniques were used to engineer deletion mutant strains [50]. A PCR ligation mutagenesis method was used to replace genes with non-polar kanamycin markers [51]. For each gene deletion, primers A and B were designed to amplify 500-600 bp upstream of the coding sequence (with ca. 50-bp overlapping the coding sequence of the gene). Primers C and D were designed to amplify 500 to 600 bp downstream of the coding sequence (with ca. 50-bp overlapping the coding sequence of the gene). Primers B and C contained *Bam*HI restriction enzyme sites for ligation of the AB and CD fragments to a non-polar kanamycin cassette digested from plasmid pALH124 [52]. Transformants were selected on BHI agar containing kanamycin. Double-crossover recombination, without introduction of nearby secondary mutations, was confirmed by PCR and Sanger sequencing using primers E and F, away from the site of recombination.

Plasmid cloning was conducted using the protein expression plasmid pBAD/His/A that contains an arabinose inducible promoter for tightly regulated protein production. PCR products for SMU_218, SMU_219 and SMU_218-219 were amplified and digested, before being ligated into digested pBAD/His/A plasmid. Correct in-frame insertion was verified with Sanger sequencing.

### Transmission electron microscopy

Overnight cultures of CRISPRi strains were diluted 1:100 into 200 μL FMC-maltose without or with 0.025% xylose, and then incubated at 37°C in a 5% CO2 incubator for 16 h. Afterwards cells were rinsed with 0.1M sodium cacodylate buffer and then fixed in 3% glutaraldehyde overnight at 4°C. The following day, cells were treated with 1.5% osmium tetroxide in the dark for 1 h at 4°C. Afterwards, the cells were mixed in equal parts with 5% agarose in PBS, collected by centrifugation at 2000 × g, and cooled to 4°C. Small chunks of the bacterial pellet plus agarose were then dehydrated in ethanol via the following steps: 30%, 50%, 70%, 80%, 90% each for 15 min, 99% 10 min, and then absolute ethanol 2 × 10 min. The dehydrated cells were embedded in an epoxy resin, sectioned and stained with uranyl acetate and lead citrate. Microscopy was conducted using a Hitachi H7600 transmission electron microscope.

### Transposon interaction screen

To look for genetic interactions of *immR*_Smu_ with other genes, a previously created Tn-seq library [4] was transformed with a *immRA*_Smu_ deletion product, described above. Transformation of the library was completed in triplicate and each transformation reaction was screened on at least ten agar plates containing spectinomycin (transposon cassette) and kanamycin (*immRA*_Smu_ deletion construct). Colonies that grew and were potentially viable *immRA*_Smu_ deletions (plus a transposon cassette) were picked and plated onto fresh antibiotic-containing media. After this, colonies were cultured overnight and frozen in glycerol at −80°C. In addition, control experiments where conducted. Controls included attempting to delete *immRA*_Smu_ with the deletion construct and selection of the Tn-seq library on kanamycin (to observe background spontaneous resistance).

### Interaction screen genome sequencing analysis

Genomic DNA was isolated from strains using a MasterPure Gram Positive DNA (Epicentre) purification kit with modifications as previously described [53]. After DNA purification, total DNA concentration and purity was measured using a NanoDrop spectrophotometer (Thermo Fisher Scientific). Purified genomic DNA was sent to the Microbial Genome Sequencing Center, and samples were sequenced according to their protocol. Sequencing reads were compared to a *S. mutans* UA159 GenBank file (NC_004350.2) using Breseq. Following genomic sequencing, transposon insertion locations were verified with PCR and Sanger sequencing (primers available in Table S6). Correct deletion of *immRA*_Smu_ was also verified using the primer pair 218_219E and 219F, followed by gel electrophoresis and comparison to expected product sizes.

### qPCR measurements of TnSmu1 excision and circularization

Quantitative PCR (qPCR) was used to calculate the excision and circularization frequency of TnSmu1. Specific primers, which only create products during excision or circularization, were used in the qPCR assay (Table S6). Primers which amplified a chromosomal single copy gene, *sloR*, close to TnSmu1 were also used to standardize results. Primers were designed using the qPCR settings in the Primer3plus online application [54]. Cell samples were grown to mid-log in rich media and DNA was collected using a MasterPure Gram Positive DNA (Epicentre) purification kit with modifications as previously described [53]. Standard curves for each primer pair were generated using eight 10-fold dilutions of PCR products, starting with 10^8^ copies/μl. Triplicates of standard curve DNAs, samples and cDNA controls were added to wells containing iQ SYBR Green Supermix (Bio-Rad) with primers (0.4 μM). Thermocycling was carried out using a CFX Touch Real-Time PCR Detection System (Bio-Rad) set to the following protocol: 40 cycles of 95°C for 10 s and 60°C for 45 s, with a starting cycle of 95°C for 30 s.

### RNA-seq analysis

RNA was extracted from OD_600_ ~0.4-0.6 bacterial cultures using the RNeasy Mini Kit. RNA was extracted from biological triplicates of CRISPRi strains carrying a short guide RNA (sgRNA) targeting SMU_218 in the presence or absence of the dCas9 inducing molecule xylose. RNA was also extracted from wild-type UA159, SMU_197c::Tn Δ*immRA*_Smu_ and SMU_201c::Tn Δ*immRA*_Smu_. Next, RNA was sent to the Microbial Genome Sequencing Center who generated RNAseq libraries and sequenced samples according to their protocol. After sequencing, read counts were aligned to the *S. mutans* UA159 genome using bioinformatics tools hosted on the Galaxy server maintained by the Research Computing Center at the University of Florida. Gene expression changes between samples were quantified with Degust (http://degust.erc.monash.edu/) using the edgeR methodology. The original RNA-seq data from this study was uploaded to the GEO database (https://www.ncbi.nlm.nih.gov/geo/) with the accession number GSE202804.

### TnSmu1 distribution bioinformatics

Upon acknowledging the TnSmu1 operon presence in *S. mutans* UA159 we isolated the gene sequences of all the individual genes by constructing a BED type file, which was parsed to the getfasta command from the Bedtools suite [55]. The resulting individual FASTA files with each gene sequence were blasted against a collection of over 600 clinical isolate genomes in our possession. The blast output was then filtered with the objective of keeping only those strains showing both a percent identity greater than 75% and a ratio between the query length and the sequence length greater than 0.9. From this filtered blast output we isolated the PID column for each isolate across all genes and generated a matrix as input for the heatmap. In parallel we conducted a Multi-locus Sequence Typing (MLST) analysis with all resulting strains in the curated blast output. The MLST sequence was then parsed to PHYML [56] to construct a Maximum-likelihood phylogeny tree. Finally the phylogenies and the heatmap were plot together in R Statistical Language [57] with packages ggplot2 [58] and GGTREE [59].

Data availability statement: the data that supports the findings of this study are available from the corresponding author. Gene expression data was uploaded to the GEO database with the accession number GSE202804.

## ACKNOWLEDGMENTS

We thank Alan Grossman at the Massachusetts Institute of Technology for helpful feedback related to this project. This work was supported by the National Institute of Dental and Craniofacial Research (NIDCR) grant DE029882 awarded to R.C.S.

