## Supplemental Material for "Activation of TnSmu1, an integrative and conjugative element, by an ImmR-like transcriptional regulator in *Streptococcus mutans*"

Shawn King[1], Allison Quick[1], Kalee King[1], Alejandro R. Walker[2], and Robert C. Shields[1*]
[1] Department of Biological Sciences, Arkansas State University, Jonesboro, Arkansas, USA
[2] Department of Oral Biology, University of Florida, Gainesville, Florida, USA

**This PDF file includes:**
Figs. S1-S5
Tables S1-S6

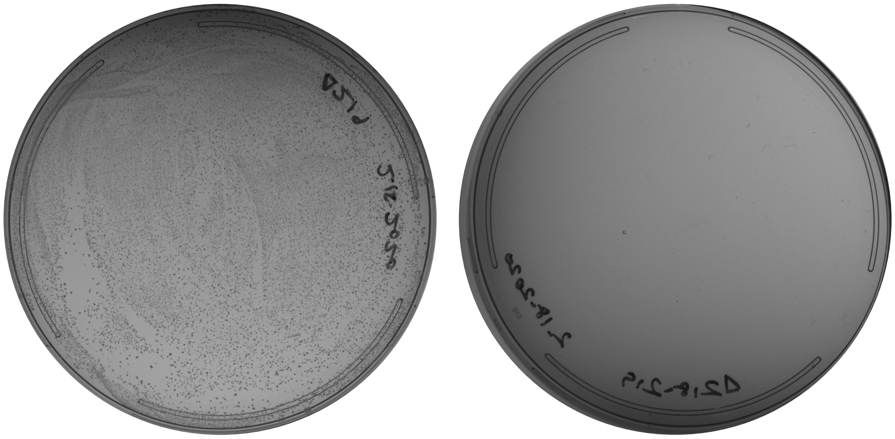

Δ219/$immA_{Smu}$ Δ218-219/$immRA_{Smu}$

**Figure S1. Colony forming units obtained from transformation experiments.** Deletion of *erfP* yielded a significant amount (lawn) of colony forming units. Under the conditions tested, deletion of *erfRP* was not permitted by *S. mutans* and zero colonies grew on transformation plates.

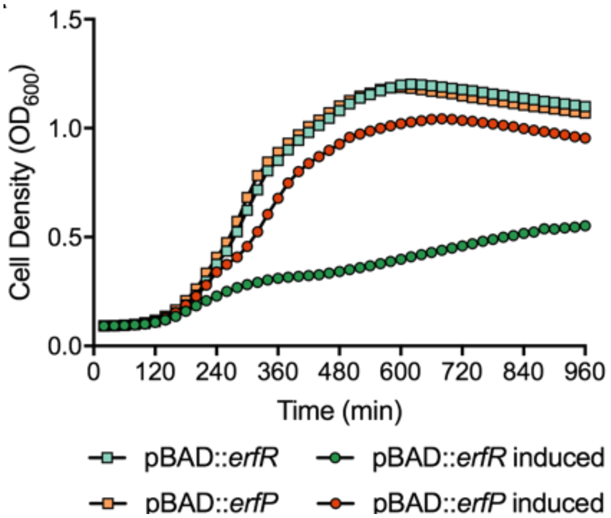


**Figure S2. Investigating the impact of *immR*$_{\mathbf{Smu}}$ and *immA*$_{\mathbf{Smu}}$ expression on *E. coli* growth.** Both *immR*$_{Smu}$ and *immA*$_{Smu}$ were cloned into an arabinose inducible protein expression system (pBAD). Induction of ImmA$_{Smu}$ had only a minor impact on *E. coli* growth (red circles). Induction of ImmR$_{Smu}$ (green circles) led to a reduction in the growth rate of *E. coli*, and the final yield after 16 h.

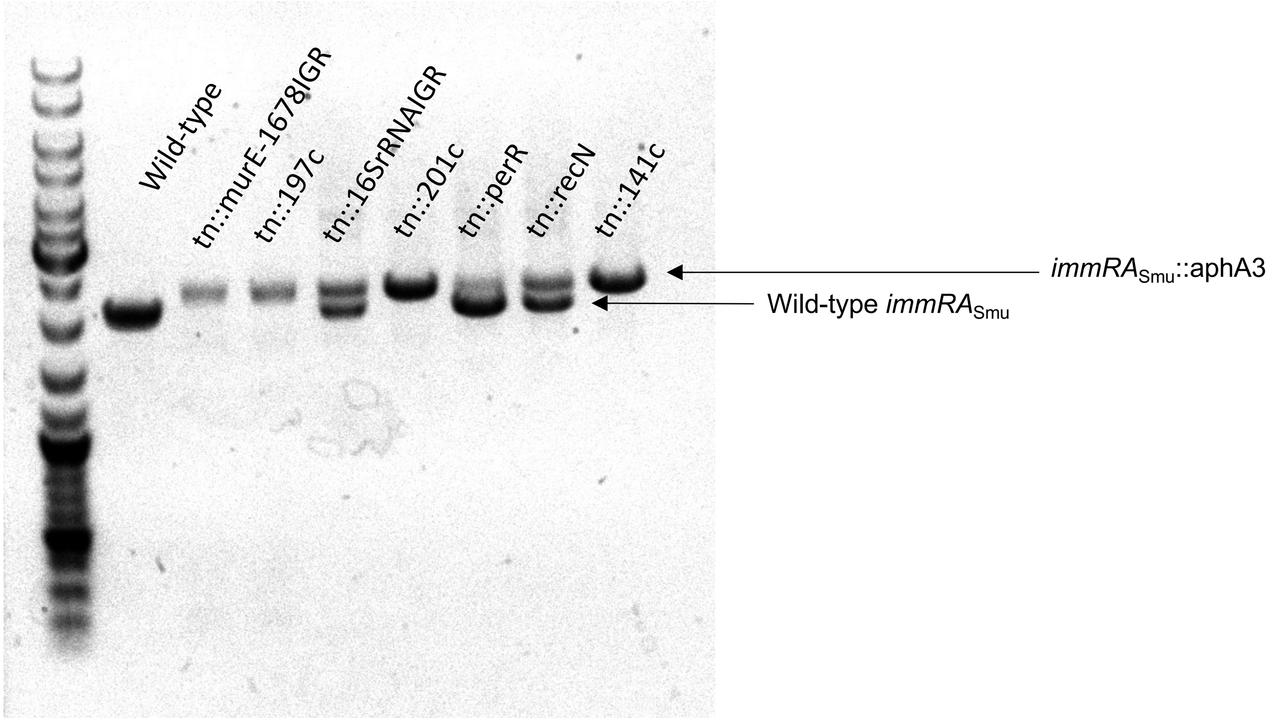


**Figure S3. PCR verification of *immRA*$_{\mathbf{Smu}}$ mutagenesis.** Correct deletion of *immRA*$_{Smu}$ (replacement with the kanamycin resistance gene *aphA3*) was confirmed with PCR. Notably there were mutants with a gene duplication event, as shown by having two PCR products of both a wild-type *immRA*$_{Smu}$ and the expected size for the mutated version.

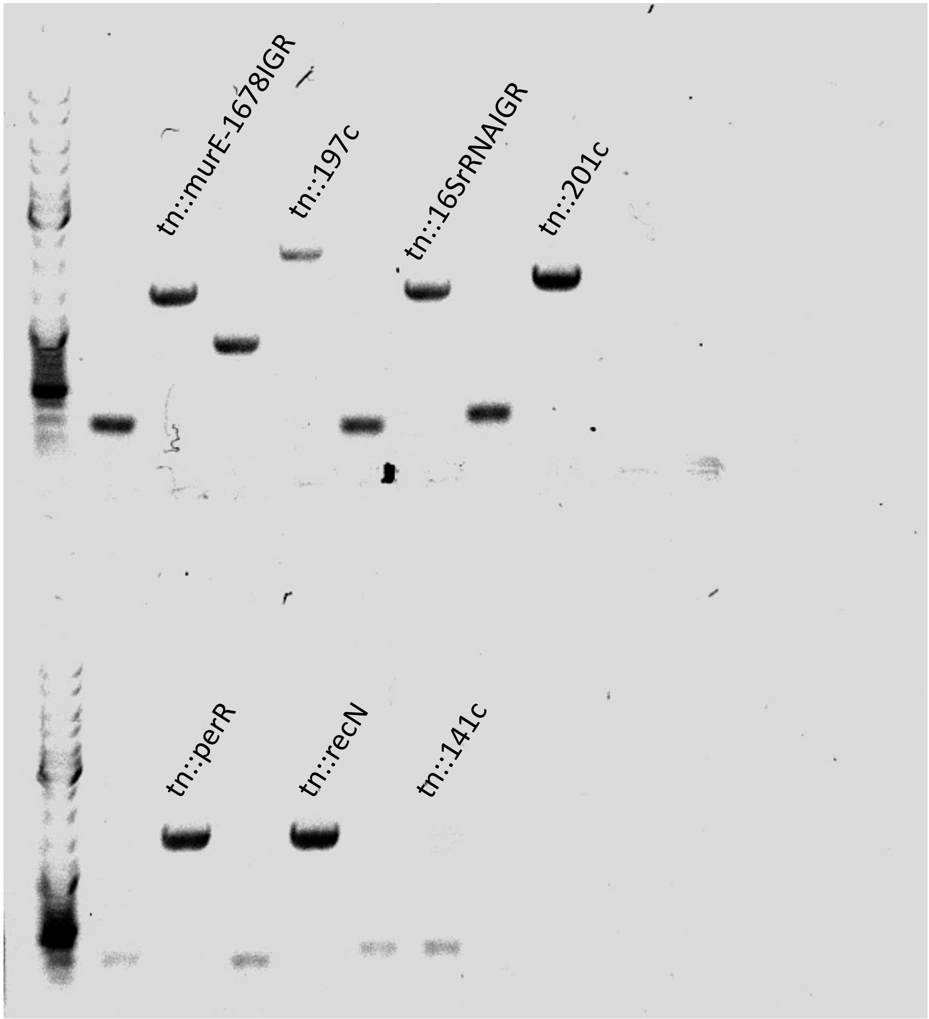


**Figure S4. PCR verification of transposon insertion.** Transposon insertions were discovered with genome sequencing and confirmed with PCR. For each expected insertion site, a primer pair was designed to amplify the region. For each mutant strain a wild-type PCR product and a PCR product from the mutant was generated and visualized with gel electrophoresis.

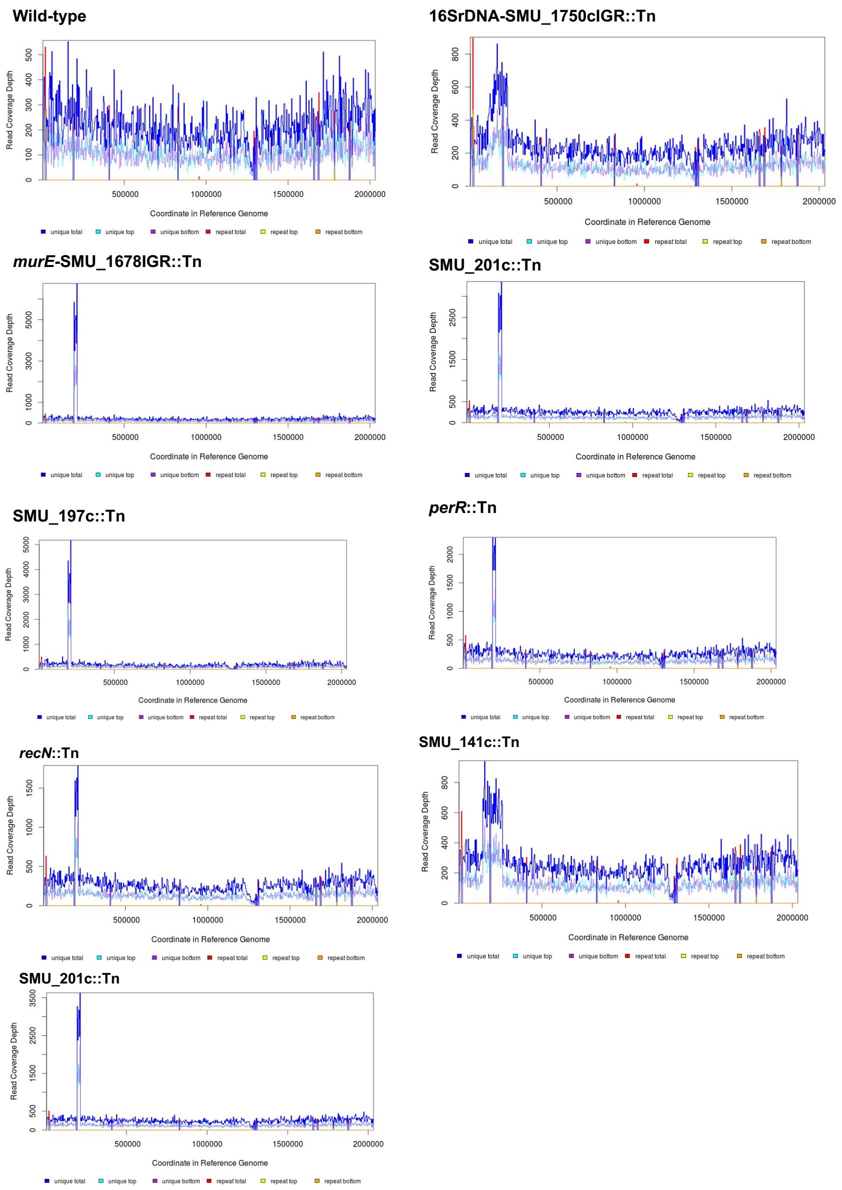


**Figure S5. Read coverage depth for sequenced genomes.** A notable trend of increased sequencing reads for the TnSmu1 region was observed for most of the double mutant strains (lacking *immRA*$_{\text{Smu}}$, or with a gene duplication event).

**Table S1. Transcriptomic changes measured by RNA-seq when *immR*$_{Smu}$ is repressed by CRISPRi**

| Gene ID | Gene name | Annotation | K number | COG | Log2 fold change | FDR |
|---|---|---|---|---|---|---|
| SMU_204c | NA | hypothetical protein | no | - | 10.03 | 3.28E-06 |
| SMU_211c | NA | hypothetical protein | no | - | 8.92 | 8.74E-06 |
| SMU_201c | NA | conserved hypothetical protein | no | - | 8.78 | 5.50E-06 |
| SMU_209c | NA | hypothetical protein | no | - | 8.76 | 3.78E-06 |
| SMU_202c | NA | conserved hypothetical protein/Streptococcus-specific protein | no | - | 8.33 | 6.02E-06 |
| SMU_197c | NA | hypothetical protein | no | - | 8.05 | 1.10E-08 |
| SMU_206c | NA | hypothetical protein | no | - | 7.97 | 3.41E-05 |
| SMU_199c | NA | hypothetical protein | no | - | 7.76 | 8.99E-06 |
| SMU_213c | NA | hypothetical protein | no | - | 7.60 | 1.69E-05 |
| SMU_210c | NA | hypothetical protein | no | - | 7.51 | 1.74E-06 |
| SMU_208c | NA | conserved hypothetical protein, FtsK/SpoIIIE family | no | - | 7.51 | 1.10E-08 |
| SMU_198c | tpn | conjugative transposon protein | no | - | 7.26 | 9.96E-08 |
| SMU_200c | NA | hypothetical protein | no | - | 7.26 | 8.87E-06 |
| SMU_205c | NA | conserved hypothetical protein | no | - | 7.06 | 1.77E-05 |
| SMU_196c | NA | immunogenic secreted protein (transfer protein) | no | - | 6.70 | 1.16E-08 |
| SMU_207c | NA | transcriptional regulator | no | - | 6.64 | 2.85E-07 |
| SMU_216c | NA | hypothetical protein | no | - | 5.66 | 2.00E-04 |
| SMU_214c | NA | hypothetical protein | no | - | 5.61 | 2.00E-04 |
| SMU_212c | NA | hypothetical protein | no | - | 5.45 | 5.04E-04 |
| SMU_221c | NA | phage-related integrase, truncated | no | - | 5.24 | 1.08E-06 |
| SMU_195c | NA | hypothetical protein | no | - | 4.83 | 4.95E-05 |
| SMU_217c | NA | conserved hypothetical protein; Streptococcus-specific protein | no | - | 4.75 | 4.37E-05 |
| SMU_215c | NA | hypothetical protein | no | - | 4.53 | 5.04E-04 |
| SMU_934 | NA | amino acid ABC transporter, permease protein | K16958 | E | 4.31 | 1.14E-07 |
| SMU_194c | NA | conserved hypothetical protein, phage-related | no | - | 4.24 | 2.00E-04 |
| SMU_962 | mmgC | acyl-CoA dehydrogenase | no | I | 4.01 | 1.24E-07 |
| SMU_219 | NA | hypothetical protein | no | - | 3.99 | 1.54E-06 |
| SMU_1753c | cas2 | CRISPR-associated endoribonuclease Cas2 | K09951 | L | 3.95 | 3.05E-07 |

| | | | | | | |
|---|---|---|---|---|---|---|
| SMU_936 | NA | amino acid ABC transporter, ATP-binding protein | K16960 | E | 3.76 | 1.49E-06 |
| SMU_220c | NA | hypothetical protein | no | - | 3.67 | 3.78E-06 |
| SMU_1155 | NA | hypothetical protein | no | - | 3.67 | 1.54E-02 |
| SMU_1757c | cas1 | CRISPR-associated protein Cas1 | no | - | 3.61 | 6.02E-06 |
| SMU_1036 | NA | conserved hypothetical protein | K01992 | V;R;C;P;G;M | 3.56 | 4.16E-05 |
| SMU_1755c | cas1 | CRISPR-associated protein Cas1 | no | - | 3.53 | 4.76E-06 |
| SMU_1889c | NA | hypothetical protein (possible relation to bacteriocin BlpU) | no | - | 3.51 | 1.67E-05 |
| SMU_193c | NA | conserved hypothetical protein | no | - | 3.49 | 5.97E-04 |
| SMU_1900 | comB | ABC transporter | K20345 | - | 3.48 | 2.99E-05 |
| SMU_961 | NA | macrophage infectivity potentiator-related protein | no | S | 3.46 | 1.18E-06 |
| SMU_218 | NA | transcriptional regulator | no | - | 3.45 | 4.43E-05 |
| SMU_653c | tauC | ABC transporter, permease protein (possible taurine transport system permease) | K02050 | P | 3.43 | 7.80E-06 |
| SMU_574c | lrg | effector of murein hydrolase | K05339 | M | 3.29 | 5.39E-03 |
| SMU_1763c | cas5 | CRISPR-associated protein, Cas5d-type | K19119 | - | 3.24 | 1.69E-05 |
| SMU_932 | NA | conserved hypothetical protein | no | T | 3.22 | 2.95E-05 |
| SMU_935 | NA | amino acid ABC transporter, permease protein | K16959 | E | 3.19 | 4.47E-06 |
| SMU_1154c | NA | conserved hypothetical protein | no | - | 3.15 | 3.15E-05 |
| SMU_1758c | cas4 | CRISPR-associated protein Cas4 | K07464 | L | 3.04 | 1.74E-06 |
| SMU_656 | mutE2 | ABC transporter, permease, possibly bacteriocin associated | K20491 | - | 2.99 | 8.42E-03 |
| SMU_933 | atmA | amino acid ABC transporter, amino acid substrate-binding protein | K16957 | E;T | 2.96 | 1.69E-05 |
| SMU_1029 | NA | hypothetical - transposon | no | - | 2.89 | 2.02E-03 |
| SMU_1156c | NA | hypothetical protein | no | - | 2.88 | 1.06E-02 |
| SMU_652c | msmK | ABC transporter, ATP-binding protein (possible nitrate transport system) | K15555 | P | 2.84 | 1.14E-04 |

| | | | | | | |
|---|---|---|---|---|---|---|
| SMU_1754c | cas1 | CRISPR-associated protein Cas1 | no | - | 2.83 | 4.62E-06 |
| SMU_59 | asl purB | adenylosuccinate lyase | K01756 | F | 2.79 | 1.34E-03 |
| SMU_1762c | csd1 | CRISPR-associated protein, Csd1-type | no | - | 2.77 | 1.22E-02 |
| SMU_1410 | frdC | fumarate reductase | K00244 | R | 2.77 | 1.31E-04 |
| SMU_1598 | celC licA | PTS system IIA component, required for cellobiose uptake and metabolism | K02759 | G | 2.74 | 2.15E-02 |
| SMU_50 | purE | phosphoribosylaminoimidazole carboxylase catalytic subunit | K01588 | F | 2.66 | 2.28E-04 |
| SMU_350 | NA | hypothetical protein | no | - | 2.61 | 8.10E-04 |
| SMU_51 | purK | phosphoribosylaminoimidazole carboxylase, ATPase subunit | K01589 | F | 2.51 | 8.52E-04 |
| SMU_1760c | csd2 | CRISPR-associated protein Csd2 | K19118 | - | 2.50 | 2.15E-05 |
| SMU_1322 | butA | acetoin reductase | K03366 | I;Q;R | 2.45 | 1.14E-04 |
| SMU_132 | hipO | amino acid amidohydrolase (hippurate amidohydrolase) | no | R | 2.41 | 1.74E-05 |
| SMU_441 | NA | transcriptional regulator | no | K | 2.38 | 2.00E-04 |
| SMU_113 | pfk | fructose-1-phosphate kinase | K00882 | G | 2.38 | 3.34E-02 |
| SMU_442 | NA | conserved hypothetical protein | no | - | 2.36 | 3.87E-04 |
| SMU_48 | purD | phosphoribosylamine-glycine ligase | K01945 | F | 2.34 | 9.19E-04 |
| SMU_1981c | comG comGF | competence protein G | K02248 | U | 2.34 | 4.77E-02 |
| SMU_145 | NA | major facilitator superfamily transporter, efflux protein | no | P | 2.32 | 6.40E-04 |
| SMU_1218 | gatA | Gln-dependent amidotransferase, subunit A | K01426 | J | 2.29 | 3.35E-05 |
| SMU_666 | argD | N-acetylornithine aminotransferase | K00821 | E | 2.28 | 2.28E-04 |
| SMU_1938c | atmE | amino acid ABC transport permease | K02072 | P | 2.27 | 5.87E-05 |
| SMU_662 | NA | conserved hypothetical protein (possible membrane protein) | K07052 | - | 2.24 | 3.46E-05 |
| SMU_94c | tpn | transposase fragment | no | - | 2.23 | 5.38E-03 |
| SMU_1970c | pheT syfB | phenylalanyl-tRNA synthetase beta subunit | K06878 | R | 2.21 | 1.94E-03 |

| | | | | | | |
|---|---|---|---|---|---|---|
| SMU_1761c | csd1 | CRISPR-associated protein, Csd1-type | K19117 | L | 2.17 | 3.37E-04 |
| SMU_1319c | NA | conserved hypothetical protein | no | S | 2.16 | 5.97E-04 |
| SMU_1578 | birA | biotin--[acetyl-CoA-carboxylase] ligase | K03524 | H | 2.14 | 8.91E-04 |
| SMU_1486c | NA | histidinol phosphatase | K04486 | E;R | 2.13 | 1.48E-04 |
| SMU_1356c | tpn | transposase | no | - | 2.12 | 3.16E-03 |
| SMU_658 | NA | conserved hypothetical protein | no | - | 2.12 | 5.75E-04 |
| SMU_1207 | fic mobC | cell filamentation / mobilization protein | K04095 | D | 2.12 | 3.21E-05 |
| SMU_2119 | opcD opuCD | ABC transport betaine/carnitine/choline permease | K05846 | E | 2.11 | 8.45E-04 |
| SMU_911c | NA | hypothetical protein | no | - | 2.10 | 2.06E-04 |
| SMU_1405c | NA | conserved hypothetical protein | K09952 | S | 2.10 | 5.85E-05 |
| SMU_1764c | cas3 | CRISPR-associated helicase | K07012 | - | 2.09 | 9.82E-05 |
| SMU_92c | | | no | - | 2.09 | 2.26E-02 |
| SMU_1770 | syv valS | valyl-tRNA synthetase | K01873 | J | 2.08 | 1.57E-03 |
| SMU_1095 | atmE pstA pstC | proline/glycine betaine ABC permease and solute binding protein | K05845 | M | 2.08 | 3.40E-04 |
| SMU_496 | cysK | cysteine synthetase A | K01738 | E | 2.07 | 1.28E-04 |
| SMU_349 | ksgA | dimethyladenosine transferase | K02528 | J | 2.07 | 5.97E-04 |
| SMU_1658 | nrgA | ammonium transporter, NrgA protein | K03320 | P | 2.07 | 4.98E-04 |
| SMU_1897 | NA | ABC transporter, ATP-binding protein; similar to BlpA | no | - | 2.06 | 8.09E-04 |
| SMU_1548c | hk11 yvfT | sensor histidine kinase | K07778 | T | 2.03 | 2.01E-04 |
| SMU_1592 | pepQ | proline dipeptidase | K01271 | E | 2.03 | 1.61E-04 |
| SMU_657 | mutG | ABC transporter, permease, possibly bacteriocin associated | K20492 | - | 2.03 | 1.54E-02 |
| SMU_872 | fruA fxpC | fructose-specific PTS system enzyme IIBC component | K02768 | G | 2.02 | 1.95E-03 |
| SMU_2038 | pttB treB | phosphotransferase system, trehalose-specific IIBC component (EIIBC-tre) | K02817 | G | 2.02 | 4.95E-05 |
| SMU_1216c | NA | ABC transporter, amino acid permease | K10009 | E | 2.00 | 9.46E-04 |

| | | | | | | |
|---|---|---|---|---|---|---|
| SMU_1203 | bcat ilvE | branched-chain amino acid aminotransferase | K00826 | E;H | 2.00 | 5.97E-04 |
| SMU_1257c | NA | conserved hypothetical protein | no | S | 2.00 | 2.47E-04 |
| SMU_1343c | pksC pksL | polyketide synthase | no | - | -2.05 | 8.48E-03 |
| SMU_2019 | rl29 rpmC | 50s ribosomal protein L29 | K02904 | J | -2.05 | 5.99E-04 |
| SMU_1059 | satC | acid tolerance protein | no | - | -2.09 | 4.01E-03 |
| SMU_748 | NA | hypothetical protein | no | - | -2.12 | 2.54E-02 |
| SMU_2018 | rpsQ rs17 | 30S ribosomal protein S17 | K02961 | J | -2.13 | 4.95E-05 |
| SMU_420 | NA | ribosomal protein L7A family | no | J | -2.13 | 5.39E-03 |
| SMU_1344c | fabD | malonyl CoA-acyl carrier protein transacylase | no | I | -2.14 | 1.59E-02 |
| SMU_1061 | ylxM | DNA-binding protein | K09787 | S | -2.17 | 7.57E-03 |
| SMU_2012 | rpsH rs8 | 30S ribosomal protein S8 | K02994 | J | -2.25 | 2.04E-04 |
| SMU_451 | NA | hypothetical protein | no | - | -2.29 | 1.46E-02 |
| SMU_758c | NA | conserved hypothetical protein | no | - | -2.32 | 3.07E-04 |
| SMU_547 | NA | conserved hypothetical protein | no | - | -2.32 | 2.47E-04 |
| SMU_276c | NA | hypothetical protein | no | - | -2.34 | 3.82E-02 |
| SMU_27 | ACP acpP | acyl carrier protein | K02078 | I;Q | -2.34 | 1.63E-03 |
| SMU_866 | NA | conserved hypothetical protein | K06960 | R | -2.35 | 4.74E-05 |
| SMU_1340 | bacA | bacitracin synthetase 1/ tyrocidin synthetase III | no | Q | -2.36 | 3.37E-04 |
| SMU_941c | NA | conserved hypothetical protein | no | - | -2.39 | 1.05E-04 |
| SMU_541 | NA | conserved hypothetical protein | no | S | -2.39 | 1.82E-02 |
| SMU_29 | purC | phosphoribosylaminoimidazole-succinocarboxamide synthase | K01923 | F | -2.47 | 3.24E-03 |
| SMU_1643c | NA | conserved hypothetical protein | no | - | -2.47 | 2.19E-02 |
| SMU_1610 | rpmG | 50S ribosomal protein L33 | K02913 | J | -2.48 | 3.06E-02 |
| SMU_1127 | rpsT | 30S ribosomal protein S20 | K02968 | J | -2.50 | 5.97E-04 |
| SMU_18 | NA | hypothetical protein | no | - | -2.51 | 3.41E-02 |
| SMU_768c | NA | conserved hypothetical protein | no | - | -2.52 | 2.25E-02 |
| SMU_1339 | bacC | bacitracin synthetase; surfactin synthetase | no | Q | -2.53 | 4.33E-04 |
| SMU_1671c | NA | conserved hypothetical protein | no | S | -2.54 | 4.52E-03 |
| SMU_1902c | NA | hypothetical protein | no | - | -2.56 | 7.13E-03 |

| | | | | | | |
|---|---|---|---|---|---|---|
| SMU_628 | holA | DNA polymerase III, delta subunit | K02340 | L | -2.65 | 1.63E-03 |
| SMU_1345c | ituA mycA | peptide synthetase similar to mycA | no | I;Q | -2.68 | 9.16E-04 |
| SMU_1507c | NA | hypothetical protein | no | - | -2.72 | 3.31E-02 |
| SMU_1907 | NA | hypothetical protein | no | - | -2.77 | 3.22E-02 |
| SMU_637c | NA | hypothetical protein | no | - | -2.78 | 4.65E-04 |
| SMU_722 | NA | hypothetical protein | no | - | -3.01 | 1.15E-02 |
| SMU_545 | NA | hypothetical protein | no | - | -3.77 | 2.73E-02 |

**Table S2. Location and analysis of *immR*$_{Smu}$ transposon double mutant strains.**

| Name | Insertion Site | Genomic Observations | Δ*erfRP** | Tn Insertion |
|---|---|---|---|---|
| **Tn-1** | IGR between *murE* and SMU_1678 | New junction SMU_191c and SMU_220c | Yes | Yes |
| **Tn-3** | SMU_197c | New junction SMU_191c and SMU_220c | Yes | Yes |
| **Tn-7** | IGR between 16S rDNA and SMU_1750c | n/a | GD | Yes |
| **Tn-8** | SMU_201c | New junction SMU_191c and SMU_220c | Yes | Yes |
| **Tn-9** | *furR* (SMU_593) | New junction SMU_191c and SMU_220c | GD | Yes |
| **Tn-10** | *recN* (SMU_585) | New junction SMU_191c and SMU_220c | GD | Yes |
| **Tn-11** | SMU_201c | New junction SMU_191c and SMU_220c | Yes | Yes |
| **Tn-12** | SMU_141 | New junction *mleS*-*sgaT* / T268T SMu.351 GTPase | Yes | Inconclusive |

*GD, gene duplication event

**Table S3. RNA-seq analysis of SMU_197c::Tn Δ*immR*$_{Smu}$ compared with *S. mutans* UA159**

| Gene ID | Gene Name | Annotation | $Log_2$ fold change | FDR |
|---|---|---|---|---|
| SMU_205c | | hypothetical protein | 8.64 | 0 |
| SMU_198c | | putative conjugative transposon protein | 8.60 | 0 |
| SMU_211c | | hypothetical protein | 8.30 | 0 |
| SMU_215c | | hypothetical protein | 8.21 | 0 |
| SMU_206c | | hypothetical protein | 8.19 | 0 |
| SMU_213c | | hypothetical protein | 8.15 | 0 |
| SMU_199c | | hypothetical protein | 8.11 | 6.665e-321 |
| SMU_204c | | hypothetical protein | 8.08 | 0 |
| SMU_212c | | hypothetical protein | 8.06 | 0 |
| SMU_217c | | hypothetical protein | 8.03 | 0 |
| SMU_216c | | hypothetical protein | 8.03 | 2.79E-288 |
| SMU_210c | | hypothetical protein | 8.00 | 0 |
| SMU_207c | | putative transposon protein | 7.95 | 0 |
| SMU_209c | | hypothetical protein | 7.95 | 0 |
| SMU_214c | | hypothetical protein | 7.95 | 3.91E-192 |
| SMU_208c | | putative transposon protein; possible DNA segregation ATPase | 7.84 | 0 |
| SMU_200c | | hypothetical protein | 7.59 | 7.37E-306 |
| SMU_202c | | hypothetical protein | 7.38 | 7.53816e-319 |
| SMU_201c | | putative transposon protein | 6.86 | 0 |
| SMU_197c | | hypothetical protein | 6.80 | 0 |
| SMU_1754c | *cas1* | CRISPR-associated protein Cas1 | 4.21 | 6.49E-178 |
| SMU_1753c | *cas2* | CRISPR-associated endoribonuclease Cas2 | 4.14 | 5.74E-296 |
| SMU_1760c | *csd2* | CRISPR-associated protein Csd2 | 4.07 | 4.56E-236 |
| SMU_1761c | *csd1* | CRISPR-associated protein, Csd1-type | 4.06 | 1.52E-305 |
| SMU_1758c | *cas4* | CRISPR-associated protein Cas4 | 4.04 | 5.92E-201 |
| SMU_1762c | *csd1* | CRISPR-associated protein, Csd1-type | 4.02 | 1.43E-206 |
| SMU_1755c | *cas1* | CRISPR-associated protein Cas1 | 3.98 | 4.82E-163 |
| SMU_1763c | *cas5* | CRISPR-associated protein, Cas5d-type | 3.96 | 3.80E-304 |
| SMU_1757c | *cas1* | CRISPR-associated protein Cas1 | 3.93 | 9.15E-173 |
| SMU_196c | | putative transfer protein | 3.89 | 1.41E-185 |
| SMU_1764c | *cas3* | CRISPR-associated helicase | 3.88 | 5.92E-293 |

| | | | | |
|---|---|---|---|---|
| SMU_194c | | conserved hypothetical protein; Bacteriophage P2 associated | 3.62 | 1.07E-107 |
| SMU_191c | | putative integrase | 3.62 | 1.31E-151 |
| SMU_1752c | | hypothetical protein | 3.61 | 3.70E-197 |
| SMU_195c | | hypothetical protein | 3.61 | 5.13E-151 |
| SMU_1750c | | hypothetical protein | 3.56 | 8.01E-124 |
| SMU_193c | | conserved hypothetical protein | 3.44 | 1.10E-79 |
| SMU_220c | | hypothetical protein | 3.35 | 4.83E-113 |
| SMU_1898 | | putative ABC transporter, ATP-binding and permease protein | 2.90 | 1.01E-95 |
| SMU_1899 | | putative ABC transporter, ATP-binding and permease protein (fragment) | 2.85 | 3.91E-29 |
| SMU_40 | | conserved hypothetical protein | 2.69 | 4.37E-12 |
| SMU_41 | | hypothetical protein | 2.52 | 1.36E-09 |
| SMU_1000 | | hypothetical protein | 2.51 | 3.67E-22 |
| SMU_1900 | *comB* | conserved hypothetical protein | 2.46 | 9.23E-49 |
| SMU_1597c | | conserved hypothetical protein | 2.18 | 2.71E-09 |
| SMU_113 | | putative fructose-1-phosphate kinase | 2.10 | 1.73E-20 |
| SMU_1598 | *celC* | putative PTS system, cellobiose-specific IIA component | 2.06 | 6.61247E-06 |
| SMU_941c | | conserved hypothetical protein | -2.07 | 1.14E-68 |
| SMU_924 | *tpx* | thiol peroxidase | -2.29 | 1.07E-32 |
| SMU_140 | *gshR* | putative glutathione reductase | -3.40 | 4.31E-32 |
| SMU_139 | *oxdC* | conserved hypothetical protein | -3.46 | 8.84E-32 |
| SMU_141 | | conserved hypothetical protein | -3.46 | 6.11E-32 |
| SMU_137 | *mleS* | malolactic enzyme | -3.68 | 4.13E-31 |
| SMU_138 | *mleP* | putative malate permease | -4.11 | 1.34E-32 |

**Table S4. RNA-seq analysis of SMU_201c::Tn Δ*immR*$_{Smu}$ compared with *S. mutans* UA159**

| Gene ID | Gene Name | Annotation | $Log_2$ fold change | FDR |
|---|---|---|---|---|
| SMU_205c | | hypothetical protein | 9.92 | 0 |
| SMU_204c | | hypothetical protein | 9.55 | 0 |
| SMU_202c | | hypothetical protein | 8.48 | 0 |
| SMU_212c | | hypothetical protein | 8.38 | 0 |
| SMU_215c | | hypothetical protein | 8.36 | 0 |
| SMU_213c | | hypothetical protein | 8.27 | 0 |
| SMU_211c | | hypothetical protein | 8.26 | 0 |
| SMU_206c | | hypothetical protein | 8.17 | 0 |
| SMU_216c | | hypothetical protein | 8.01 | 6.43E-287 |
| SMU_214c | | hypothetical protein | 8.00 | 6.13E-194 |
| SMU_217c | | hypothetical protein | 7.97 | 0 |
| SMU_209c | | hypothetical protein | 7.95 | 0 |
| SMU_210c | | hypothetical protein | 7.85 | 0 |
| SMU_208c | | putative transposon protein; possible DNA segregation ATPase | 7.64 | 0 |
| SMU_201c | | putative transposon protein | 7.60 | 0 |
| SMU_207c | | putative transposon protein | 7.45 | 0 |
| SMU_197c | | hypothetical protein | 4.80 | 6.03E-253 |
| SMU_198c | | putative conjugative transposon protein | 4.75 | 0 |
| SMU_195c | | hypothetical protein | 4.70 | 3.51E-242 |
| SMU_200c | | hypothetical protein | 4.70 | 7.29E-143 |
| SMU_196c | | putative transfer protein | 4.69 | 9.50E-256 |
| SMU_191c | | putative integrase | 4.65 | 1.54E-232 |
| SMU_199c | | hypothetical protein | 4.50 | 2.24E-129 |
| SMU_194c | | conserved hypothetical protein; Bacteriophage P2 associated | 4.49 | 2.77E-158 |
| SMU_193c | | conserved hypothetical protein | 4.17 | 3.00E-113 |
| SMU_1753c | *cas2* | CRISPR-associated endoribonuclease Cas2 | 3.91 | 8.56E-269 |
| SMU_1754c | *cas1* | CRISPR-associated protein Cas1 | 3.88 | 6.10E-155 |
| SMU_1757c | *cas1* | CRISPR-associated protein Cas1 | 3.79 | 1.61E-162 |
| SMU_1755c | *cas1* | CRISPR-associated protein Cas1 | 3.76 | 1.96E-147 |

| | | | | |
|---|---|---|---|---|
| SMU_1762c | *csd1* | CRISPR-associated protein, Csd1-type | 3.75 | 1.57E-183 |
| SMU_1761c | *csd1* | CRISPR-associated protein, Csd1-type | 3.75 | 3.91E-267 |
| SMU_1763c | *cas5* | CRISPR-associated protein, Cas5d-type | 3.72 | 7.08E-273 |
| SMU_1758c | *cas4* | CRISPR-associated protein Cas4 | 3.71 | 1.04E-173 |
| SMU_1760c | *csd2* | CRISPR-associated protein Csd2 | 3.70 | 2.56E-200 |
| SMU_1764c | *cas3* | CRISPR-associated helicase | 3.69 | 1.54E-267 |
| SMU_1752c | | hypothetical protein | 3.56 | 3.43E-192 |
| SMU_1750c | | hypothetical protein | 3.40 | 3.66E-114 |
| SMU_220c | | hypothetical protein | 3.15 | 3.13E-101 |
| SMU_40 | | conserved hypothetical protein | 2.65 | 1.25E-11 |
| SMU_1029 | | conserved hypothetical protein | 2.55 | 5.63E-17 |
| SMU_41 | | hypothetical protein | 2.41 | 1.13E-08 |
| SMU_1899 | | putative ABC transporter, ATP-binding and permease protein (fragment) | 2.29 | 9.73E-19 |
| SMU_1898 | | putative ABC transporter, ATP-binding and permease protein | 2.18 | 1.37E-55 |
| SMU_1539 | | putative 1,4-alpha-glucan branching enzyme | -2.19 | 4.59E-58 |
| SMU_1538 | | putative glucose-1-phosphate adenylyltransferase; ADP-glucose pyrophosphorylase | -2.26 | 1.36E-63 |
| SMU_141 | | conserved hypothetical protein | -3.55 | 3.30E-33 |
| SMU_140 | *gshR* | putative glutathione reductase | -3.79 | 4.47E-38 |
| SMU_139 | *oxdC* | conserved hypothetical protein | -3.82 | 3.70E-37 |
| SMU_137 | *mleS* | malolactic enzyme | -4.18 | 8.59E-38 |
| SMU_138 | *mleP* | putative malate permease | -4.44 | 1.24E-36 |

**Table S5. Strains and plasmids used in this study.**

| Strain | Description | Source |
|---|---|---|
| *S. mutans* strains | | |
| UA159 | Wild-type | Burne Lab |
| Δ*immA*$_{Smu}$ | SMU_219::*aphA3* | This work |
| Tn-1 | IGRmurE_SMU_1678::Tn Δ*immRA*$_{Smu}$::*aphA3* | This work |
| Tn-3 | SMU_197c::Tn *immRA*$_{Smu}$::*aphA3* | This work |
| Tn-7 | IGR16SrDNA_SMU_1750c::Tn Δ*immRA*$_{Smu}$::*aphA3* | This work |
| Tn-8 | SMU_201c::Tn Δ*immRA*$_{Smu}$::*aphA3* | This work |
| Tn-9 | SMU_593::Tn Δ*immRA*$_{Smu}$::*aphA3* | This work |
| Tn-10 | SMU_585::Tn Δ*immRA*$_{Smu}$::*aphA3* | This work |
| Tn-11 | SMU_201c::Tn Δ*immRA*$_{Smu}$::*aphA3* | This work |
| Tn-12 | SMU_141::Tn Δ*immRA*$_{Smu}$::*aphA3* | This work |
| CRISPRi sgRNA-*lacG* | pDL278::P$_{xyl}$-*dcas9* pPM::sgRNA-*lacG* | (1) |
| CRISPRi sgRNA-*immR*$_{Smu}$ | pDL278::P$_{xyl}$-*dcas9* pPM::sgRNA-*immR*$_{Smu}$ | (1) |
| *E. coli* strains | | |
| 10-beta | Cloning host, derivative of DH10B | New England Biolabs |
| BL21(DE3) | Cloning host suitable for protein expression | New England Biolabs |
| Plasmids | | |
| pBAD/His/A | Protein expression vector with an *araBAD* promoter for tightly regulated expression | Thermo Fisher Scientific |

**Table S6. Oligonucleotides used in this study.**

| Name | Oligonucleotide sequence (5'-3') | Description |
|---|---|---|
| *S. mutans cloning* | | |
| 219A | TGTTCCCAGAACGCTTAAAA | Deletion of SMU_219 |
| 219B | ATGCGGATCCTGATGGTAACTCCAAGTTTTTCTG | Deletion of SMU_219 |
| 219C | ATGCGGATCCATGAAAACATTGTTAAAGAAGAAGC | Deletion of SMU_219 |
| 219D | TGATGAAGGCAAATTGTGGA | Deletion of SMU_219 |
| 219E | TGAGGTTAACTAGATTTTTAACAG | Sequencing deletion of SMU_219 |
| 219F | GGAACCTATTATGTTTATAAGTG | Sequencing deletion of SMU_219 |
| 218_219A | TCTAGCATGCGCTCATATCCT | Double deletion of SMU_218 and SMU_219 |
| 218_219B | ATGCGGATCCTGCTTCTAGGCGCAGAGATT | Double deletion of SMU_218 and SMU_219 |
| 218_219E | ACCACTCATCTGCTTAATAA | Sequencing double deletion of SMU_218 and SMU_219 |
| *E. coli cloning* | | |
| pBAD218Fv2 | GATCGGTACCATATGTTCCCAGAACGCTTA | SMU_218 cloning into pBAD/His/A |
| pBAD218R | GATCGAATTCCGTTTAGTTATTTTTCTGATTTTTGA | SMU_218 cloning into pBAD/His/A |
| pBAD219Fv2 | GATCGGTACCATATGAACTTATCAAAAATTGTTAGAGA | SMU_219 cloning into pBAD/His/A |
| pBAD219R | GATCGAATTCGTATCTTTTTATTTTGGTTTTGTCA | SMU_219 cloning into pBAD/His/A |
| pBADseqF | ATGCCATAGCATTTTTATCC | pBAD/His/A sequencing |
| pBADseqR | GATTTAATCTGTATCAGG | pBAD/His/A sequencing |
| *Transposon verification* | | |
| Tn-3_Ins_SeqF | GCGATCACAGAAGCACAAAA | Interaction screen transposon insertion verification |
| Tn-3_Ins_SeqR | GACAGAGACAAGGCCCAAAA | Interaction screen transposon insertion verification |
| Tn-1_Ins_SeqF | CGCCAACAGTCGTTAAGGTT | Interaction screen transposon insertion verification |
| Tn-1_Ins_SeqR | CAGGAGTGTTAGGGATTCCTTG | Interaction screen transposon insertion verification |
| Tn-7_Ins_SeqF | GACCGAAACTTAGGCTTGGA | Interaction screen transposon insertion verification |
| Tn-7_Ins_SeqR | TAAAACCCAAGGACGGACTG | Interaction screen transposon insertion verification |
| Tn-8_Ins_SeqF | AAAGGGACAGGAATCTAGGG | Interaction screen transposon insertion verification |
| Tn-8_Ins_SeqR | CCTGAAAAAGGAGCCATCTG | Interaction screen transposon insertion verification |

| | | |
|---|---|---|
| Tn-9_Ins_SeqF | TCCTGATTGATGAAGGCTTTG | Interaction screen transposon insertion verification |
| Tn-9_Ins_SeqR | TGACTCACCTCCTATTTCCCATA | Interaction screen transposon insertion verification |
| Tn-10_Ins_SeqF | CGGTAAATGACCTCGCTTTT | Interaction screen transposon insertion verification |
| Tn-10_Ins_SeqR | TGAGACGAGACAGCTCTCCA | Interaction screen transposon insertion verification |
| Tn-12_Ins_SeqF | GGCTGCTCAGACAGGAAATC | Interaction screen transposon insertion verification |
| Tn-12_Ins_SeqR | CTCACACCAAGCAATGATGG | Interaction screen transposon insertion verification |
| *TnSmu1 qPCR* | | |
| CircTnSmu1.1F | AAATTTTCTCCCAAAAATTATCAAA | qPCR for circular TnSmu1 |
| CircTnSmu1.1R | AAAGAGTTTAAAGAGGTTGAACAAA | qPCR for circular TnSmu1 |
| ExcisTnSmu1F | CAATTCCTACTGCCCGTGTT | qPCR for TnSmu1 excision |
| ExcisTnSmu1R | GATATTTGGGCGGTTGCTAA | qPCR for TnSmu1 excision |
| TnSmu1sloRF | TTTATCGCAAGCATCGTCTG | qPCR for chromosomal gene |
| TnSmu1sloRR | GGCTGTCCATGTTGAGGAAT | qPCR for chromosomal gene |
